# Dicloxacillin and flucloxacillin inhibit hepatic uptake transporters – in vitro investigations and physiologically based pharmacokinetic modelling

**DOI:** 10.1101/2025.10.21.683780

**Authors:** Noora Sjöstedt, Ogochukwu U Amaeze, Jeroen JMW van den Heuvel, Tore B Stage, Jan B Koenderink, Nina Isoherranen, Heidi Kidron, Erkka Järvinen

**Affiliations:** Division of Pharmaceutical Biosciences, Faculty of Pharmacy, University of Helsinki, Helsinki, Finland; Department of Pharmaceutics, School of Pharmacy, University of Washington, Seattle, WA United States; Department of Pharmacy, Pharmacology and Toxicology, Radboud University Medical Center, Nijmegen, Netherlands; Clinical Pharmacology, Pharmacy, and Environmental Medicine, Department of Public Health, University of Southern Denmark, Odense, Denmark; Department of Clinical Pharmacy and Biopharmacy, Faculty of Pharmacy, University of Lagos, Lagos, Nigeria; Institute of Biotechnology, Helsinki Institute of Life Science, University of Helsinki, Helsinki, Finland

**Author notes:** Corresponding author: Erkka Järvinen. CONFLICT OF INTEREST: *T.B.S. has given paid lectures for Pfizer and Eisai, consulted for Pfizer, been an expert witness for Sandoz A/S, and collaborated with Novo Nordisk A/S, all unrelated to the work reported in the present article. N.I. reports consultancy agreements with Merck and Boehringer-Ingelheim, and honoraria from ASPET, Elsevier and the US National Institutes of Health. Other authors declare no conflict of interest.

## Abstract

Dicloxacillin and flucloxacillin are β-lactamase-resistant penicillin antibiotics that have been in clinical use for over 50 years. While both antibiotics are known to induce cytochrome P450 enzymes, there is limited information available regarding their interactions with drug transporters. Here, we investigated the in vitro transport and inhibition of hepatic organic anion transporting polypeptides (OATPs) and renal organic anion transporters (OATs) by these antibiotics in recombinant transporter overexpressing HEK293 cells. We also investigated the transport of these antibiotics by efflux transporters, as well as their inhibition of breast cancer resistance protein (BCRP) and P-glycoprotein (P-gp) using a HEK293 membrane vesicle transport assay. Dicloxacillin and flucloxacillin inhibited rosuvastatin transport by OATP1B1, OATP1B3 and OATP2B1, and the inhibition was strongest for OATP1Bs with IC_50_ values of 3.9 µM and 31 µM (OATP1B1) and 6.7 µM and 21 µM (OATP1B3) for dicloxacillin and flucloxacillin, respectively. Both antibiotics also inhibited BCRP-mediated rosuvastatin transport with IC_50_ values of 166 µM (dicloxacillin) and 379 µM (flucloxacillin), while P-gp-mediated transport of N-methyl-quinidine was inhibited to a lesser extent. All OATPs and OATs transported dicloxacillin and flucloxacillin. Static model predictions indicated that the inhibition of OATPs, BCRP and P-glycoprotein by both compounds may be clinically relevant. We further developed and verified physiologically based pharmacokinetic (PBPK) models for dicloxacillin and flucloxacillin. PBPK model simulations predicted no major change in rosuvastatin, a substrate for OATPs and BCRP, pharmacokinetics, when co-administered with dicloxacillin or flucloxacillin. Simulations with dicloxacillin and P-gp substrates dabigatran or digoxin also predicted limited inhibition of P-gp transport.

## INTRODUCTION

Dicloxacillin and flucloxacillin are β-lactamase resistant penicillin antibiotics used to treat staphylococcal infections, such as skin and soft tissue infections, but also endocarditis.^1^ In Europe, flucloxacillin is among the top four antibiotics consumed.^2^ Pharmacokinetics of both antibiotics are characterized by rapid clearance, high plasma protein binding, minimal metabolism and excretion via urine.^3–9^ Because of the short elimination half-lives of dicloxacillin and flucloxacillin, they are administered three to four times per day.^1,8,10^

Dicloxacillin and flucloxacillin were only recently found to induce cytochrome P450 enzymes (CYPs) in clinical trials and in vitro.^8,10,11^ Moreover, the reduction in warfarin efficacy when dicloxacillin or flucloxacillin therapy is initiated, suggests a pharmacokinetic interaction between these antibiotics and warfarin.^11^ Dicloxacillin weakly induces intestinal P-glycoprotein (P-gp) in humans and in vitro but this does not translate to decreased efficacy of direct oral anticoagulants that are P-gp substrates.^12^ Only a few drugs are known to affect the pharmacokinetics of dicloxacillin and flucloxacillin. Probenecid slightly increases the plasma concentration of dicloxacillin and flucloxacillin, while cyclosporine increases and rifampicin slightly decreases the plasma concentration of dicloxacillin.^6,7,9^

Drug transporters involved in the disposition of dicloxacillin and flucloxacillin are poorly characterized. As negatively charged carboxylic acids with a molecular weight of 450-470 Da, dicloxacillin and flucloxacillin are likely to interact with hepatic organic anion transporting polypeptides (OATPs) and renal organic anion transporters (OATs).^13,14^ More importantly, in vitro reports suggest that these drugs may strongly inhibit human OATP1B1 and OATP1B3, and dicloxacillin may inhibit OAT3.^15,16^ Similarly, knowledge on efflux transporters involved in the transport of dicloxacillin and flucloxacillin or efflux transporters inhibited by these drugs is limited. Dicloxacillin was identified as a P-gp substrate in a cell assay, while it did not inhibit multidrug resistance-associated proteins (MRP) 2, 3 or 4 in membrane vesicle assays.^17,18^ On the other hand, flucloxacillin did not inhibit breast cancer resistance protein (BCRP) or MRP2 in a cell assay.^19^

In this study, we comprehensively characterized the in vitro transport and inhibition of human OATP1B1, OATP1B3 and OATP2B1 and OAT1-OAT4 by dicloxacillin and flucloxacillin. We also investigated their transport by human efflux transporters and studied the inhibition of intestinal efflux transporters BCRP and P-gp. Finally, we developed physiologically based pharmacokinetic (PBPK) models for both antibiotics to translate in vitro inhibition data into human drug-drug interaction (DDI) predictions.

## METHODS

### Chemicals and reagents

Flucloxacillin and fluorescein sodium salt, dicloxacillin sodium salt monohydrate, sodium bicarbonate, sodium butyrate, HEPES, and analytical grade solvents methanol, acetonitrile, phosphoric acid and formic acid were acquired from Sigma-Aldrich (St. Louis, MO, USA). Rosuvastatin calcium salt and rosuvastin-d_6_ sodium salt were from Toronto Research Chemicals (North York, Ontario, Canada). N-methyl-quinidine was from Solvo Biotechnology (Szeged, Hungary). Tritium-labelled methotrexate disodium salt was from Moravek (Brea, CA, USA). Dulbecco’s Modified Eagle Medium (DMEM) supplemented with high glucose, GlutaMax, HEPES, phenol red and without sodium pyruvate, referred hereinafter as DMEM, Hank’s Balanced Salt Solution with calcium, magnesium, glucose and without sodium bicarbonate and phenol red, poly-D-lysine solution, fetal bovine serum, Nunc cell-culture treated 48-well plates, and Corning BioCoat poly-D-lysine 24-well plates were from ThermoFisher Scientific (Waltham, MA, USA).

### Transient expression of transporters in HEK293 cells

The human transporters OATP1B1, OATP1B3, OATP2B1 and OAT1 to OAT4 and a negative control protein (enhanced yellow fluorescent protein, eYFP) were transiently expressed in human embryonic kidney 293 (HEK293) cells by recombinant baculovirus-mediated transduction. Baculoviruses carrying human genes for the transporters were prepared as reported previously.^20–22^ HEK293 cells were cultured in DMEM containing 10% fetal bovine serum at 37 °C and 5% CO_2_ and passaged two to three times per week. For the transport studies, 0.05×10^6^ cells for 48-well plates (OATPs) or 0.06×10^6^ cells for 24-well plates (OATs) were seeded onto cell-culture treated well plates coated with poly-D-lysine. After one day, cells were transduced to produce recombinant proteins by changing media to DMEM containing 30 µl (48-well plates) or 60 µl (24-well plates) of recombinant baculovirus and 3 mM (OATs) or 5 mM (OATPs) sodium butyrate in a total media volume of 500 µl. The cells were cultured for a further two days before the transport assays.

### Transport assays in HEK293 cells overexpressing OATP and OAT transporters

For the transport studies, media were removed and cells were washed once with warm transport uptake buffer (TP-buffer – pH 7.4 HBSS with 25 mM HEPES and 4 mM sodium bicarbonate). Each experiment included triplicate (OATP) or duplicate (OAT) samples for every test condition. The uptake was started by removing liquid from wells and applying warm TP-buffer (200 µl for 24-well plates or 125 µl for 48-well plates) containing a test compound to cells. Transport assays were terminated after a pre-specified incubation time (reported in figure legends) by removing substrate solution and washing cells three times with cold TP-buffer (300 µl for 48-well plates and 400 µl for 24-well plates). For the liquid chromatography mass spectrometry (LC-MS) analysis, compounds were extracted from cells with 125 µl (48-well plates) or 250 µl (24-well plates) of an ultrapure-water solution containing 75% methanol and 0.1% formic acid to wells and shaking at 250 RPM for 30 min. For LC-MS analysis, internal standards (dicloxacillin for flucloxacillin, flucloxacillin for dicloxacillin and rosuvastatin-d_6_ for rosuvastatin) were spiked in the extraction solution before applying it to cells. Samples were transferred to well plates, centrifuged at 3000 g for 30 min and transferred to a new well plate for the analysis. All substrate and inhibitor stocks were prepared in dimethyl sulfoxide (DMSO), and the DMSO concentration did not exceed 0.3% in the transport assays.

### Inhibition assays in HEK293 cells overexpressing OATP and OAT transporters

In the OATP inhibition assays, inhibitors in warm TP-buffer were directly applied after media removal, followed by a 30-minute preincubation step in the presence of inhibitors. No preincubation step was included in the OAT inhibition assays. The inhibition assays were conducted as described above, except that inhibitors were included in the substrate solution. The rosuvastatin concentration was 2.5 µM in the OATP inhibition assays, and the transport assays were conducted for 2 min.^23^ For inhibition studies of OAT1 and OAT3, 5 µM fluorescein was assayed within 5 min, while 34 nM methotrexate within a 10 min assay was employed as the substrate for OAT2 and OAT4.^24–27^ OAT1 and OAT3 cells were lysed with 200 µl of 1 M sodium hydroxide for 10 min, and 150 µl of these samples were transferred to a black well plate for fluorescein quantification. OAT2 and OAT4 cells were lysed with 200 µl of 0.1% Triton-X100 for 10 min, and 150 µl of the samples were mixed with 4 ml of scintillation cocktail (Opti-Fluor from PerkinElmer, Waltham, MA, USA). Each inhibition assay included a vehicle control without an inhibitor.

### Efflux transport and inhibition assays in HEK293 membrane vesicles

Membrane vesicles, 5 µl (7.5 µg of total protein), prepared as previously reported ^22^, were aliquoted to wells in a conical-shaped 96-well plate. The plate was incubated for 10 min at +37 °C before the initiation of transport by adding 25 µl of TS-buffer (10 mM Tris, 250 mM Sucrose, pH adjusted to 7.4 with HEPES) containing magnesium chloride (10 mM final in the reaction), adenosine triphosphate (ATP) or adenosine monophosphate (AMP), 4 mM final in the reaction, and the transporter substrate and inhibitor. Each experiment included triplicate samples for both ATP and AMP samples. The concentrations of rosuvastatin and N-methyl-quinidine were 1 µM in the BCRP and P-gp vesicle inhibition assays, respectively.^28,29^ The transport reaction times were 1 and 2 min for BCRP and P-gp, respectively.^28,29^ The vehicle concentration (DMSO) was 1.1% in the efflux transporter inhibition assays. The transport reactions were quenched by adding 200 µl of cold TS-buffer to each well and transferring the samples to a 96-multiwell 1.0 μm glass fiber filter plate (Merck Millipore, Darmstadt, Germany) kept under vacuum. Wells were further washed four times with cold TS-buffer, after which the plate was dried under vacuum. The compounds were eluted from the plate using 100 µl of 75% methanol containing 0.1% formic acid to each well and incubating at 250 RPM for 30 min before centrifing the samples to an analysis plate at 2800 g for 2 min. For the rosuvastatin samples, the elution solvent contained rosuvastin-d_6_ as the internal standard.

### Analytical methods

An LC-MS system consisting of Acquity I class UPLC connected to triple quadrupole Xevo TQ-S mass spectrometer equipped with an electrospray ionization source, from Waters (Milford, MA, USA), was employed to quantify dicloxacillin, flucloxacillin and rosuvastatin. N-methyl-quinidine was quantified with Agilent Technologies (Santra Clara, CA, USA) 1100 HPLC system equipped with a fluorescence detector. Each batch of LC-MS runs included a standard curve, and quantification was based on the analyte to internal standard peak area ratio plotted against the analyte concentration. An external standard curve calibration was employed for N-methyl-quinidine. Details of LC-MS and HPLC methods are reported in *Supplementary methods*. Fluorescein was quantified with a fluorescence plate reader (Victor X3) by employing a standard curve and wavelengths 485 nm (excitation) and 535 nm (emission). An automatic liquid scintillation counter (Hidex 600 SL from Hidex, Turku Finland) was employed for quantification of tritium-labelled methotrexate.

### PBPK modelling

Full-PBPK models for dicloxacillin and flucloxacillin were developed in Simcyp version 23 (Certara UK Limited, Sheffield, UK). Further details of models are reported in *Supplementary methods and results* (Table S2 and S3). Published clinical pharmacokinetic data were divided into two datasets, one for model development and the other for model verification (Table S4 and S5). For the DDI simulations, Simcyp built-in library models for rosuvastatin, dabigatran etexilate and digoxin were employed.

### Data analysis and reporting

For the inhibition data analysis, the average of measured concentrations in the control cells was subtracted from the average of measured concentrations in transporter-expressing cells for each inhibitor concentration and then normalized to the vehicle sample that represents 100% transport activity. Equation 1 was fit to the inhibition data. In Equation 1, Inhibition represents the percent inhibition of transporter-mediated uptake at an inhibitor concentration of I and IC_50_ is the half-maximal inhibitory concentration.

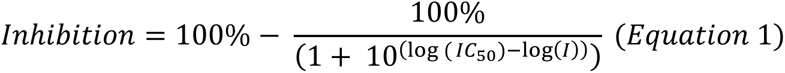

Equation 2 (the Michaelis-Menten equation) was fit to the transport kinetic assay data. In Equation 2, Uptake is the rate of uptake in transporter-expressing cells from which the uptake in the control cells is subtracted at the concentration S of dicloxacillin or flucloxacillin, V_max_ is the maximum rate of transport, and K_m_ is the Michaelis constant.

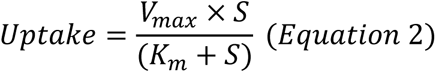

The nonlinear least squares method (nls) in the R programming language environment (R version 4.3.1) was employed for fitting Equations 1 and 2 to the data.

To evaluate potential clinical DDIs for transporters, FDA R values were calculated according to the FDA In Vitro Drug Interaction Studies guidance.^30^ Equation 3 was used to estimate the maximum unbound plasma inhibitor concentration at the inlet to the liver (I_in,max_) for OATPs, where f_u,p_ is the unbound fraction in plasma, C_max_ is the maximal total concentration in plasma, F_a_ is the fraction absorbed after oral administration, F_g_ is the fraction available after intestinal metabolism, k_a_ is the first order absorption rate constant in vivo, dose is the oral dose, Q_h_ is the hepatic blood flow and R_B_ is the blood-to-plasma concentration ratio.

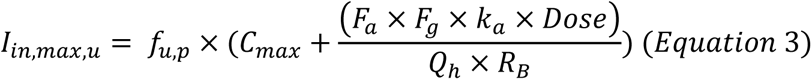

The values of F_a_*F_g_ and k_a_ were set to 1 and 0.1/min to present a worst-case estimation according to FDA’s recommendation, since no experimental bioavailability data is available for dicloxacillin or flucloxacillin.^30^ Hepatic blood flow (Q_h_) was set to 1.620 l/min.^30^ A typical oral dose for dicloxacillin and flucloxacillin is 1000 mg. Experimental R_B_ is 0.65 for flucloxacillin and was assumed to be same for dicloxacillin.^31^ The unbound fraction in plasma for dicloxacillin and flucloxacillin were set to 0.02 and 0.06, respectively.^4^ The total C_max_ values for dicloxacillin and flucloxacillin at 1000 mg dose are reported as 67 µM and 46 µM, respectively.^6,8^ The resulting calculated I_n,max,u_ values for dicloxacillin and flucloxacillin were 5.51 µM and 15.7 µM, respectively. For OATs, the maximal unbound plasma concentration of the interacting drug at steady state (I_max,u_)^30^ was calculated by C_max*_f_u_ resulting in 1.34 µM and 2.76 µM for dicloxacillin and flucloxacillin, respectively. The intestinal luminal concentration (I_gut_)^30^ was estimated by dose/250 ml, resulting in 8505 µM and 8813 µM for dicloxacillin and flucloxacillin, respectively.

## RESULTS

### Transporter inhibition studies

Dicloxacillin and flucloxacillin inhibited rosuvastatin transport into HEK293 cells expressing the human hepatic OATPs (Figure 1). We pre-incubated the cells for 30 min in the presence of inhibitors, since this procedure is currently recommended for OATPs to fully reveal the strength of inhibition.^30,32,33^ The IC_50_ values were lower for dicloxacillin than flucloxacillin (Table 1). We replicated the inhibition experiment by employing fluorescent substrates for OATPs (Figure S1) and the obtained IC_50_ values (Table S6) were comparable to the experiments with rosuvastatin as the OATP substrate (Table 1). We also tested whether the inhibition of OATP1B1 by dicloxacillin and flucloxacillin is reversible (Figure S2). We were able to wash out most of the inhibition, which indicates that the inhibition of OATP1B1 is reversible and is not caused by covalent binding of dicloxacillin and flucloxacillin to OATPs, as reported previously for these compounds with human serum albumin.^34^ Moreover, we determined the K_i_-value for the inhibition of OATP1B1 by dicloxacillin, since this was the strongest observed inhibition. The experimental K_i_-value for the competitive inhibition of OATP1B1 by dicloxacillin was 3.65 µM (2.33-5.71, 95% CI), which is close to the experimental IC_50_-value (Figure S3, Table 1).

**Figure 1.**
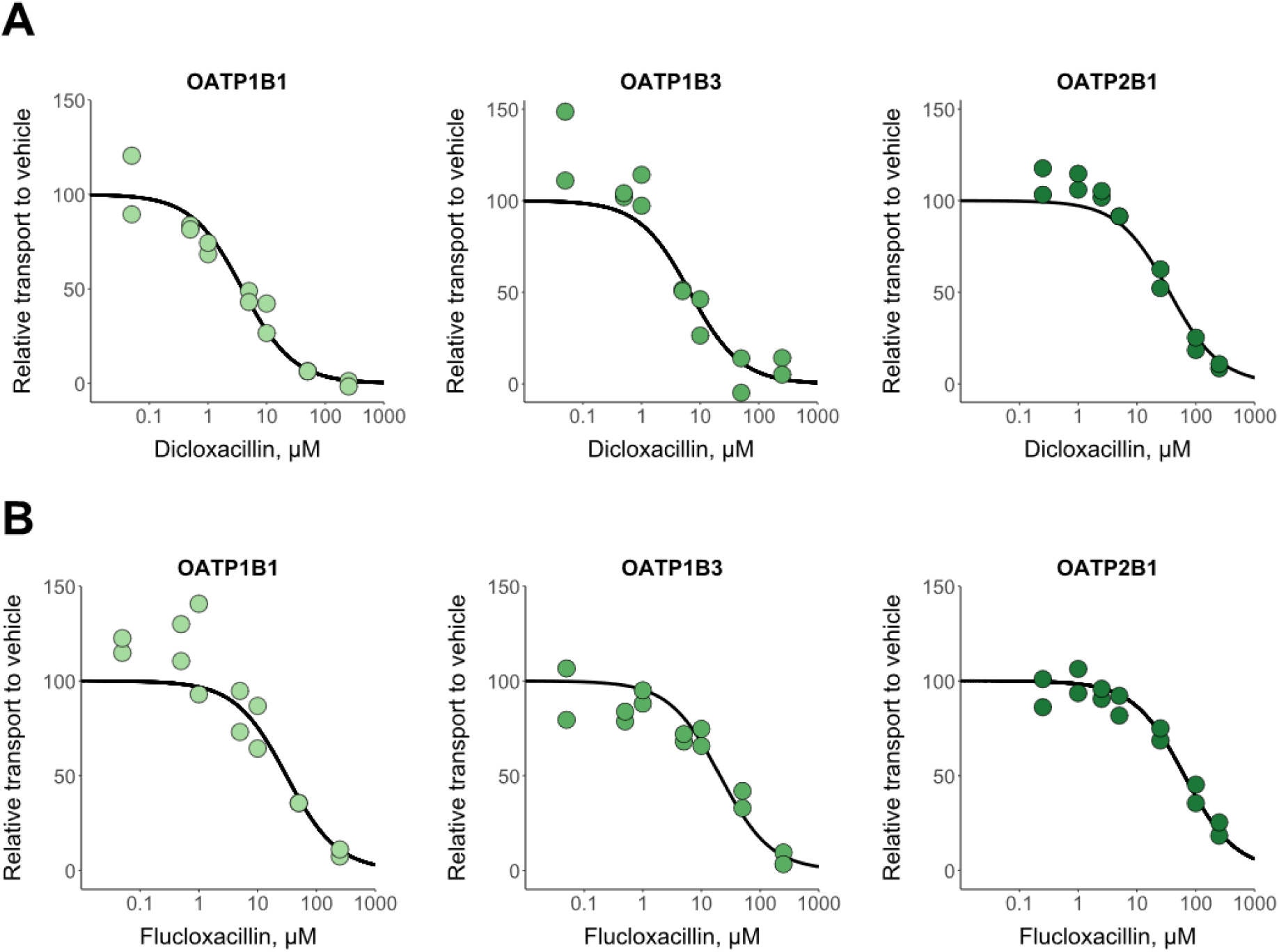
In vitro inhibition of OATPs by dicloxacillin (A) and flucloxacillin (B). Rosuvastatin transport into OATP1B1-, OATP1B3-, OATP2B1- and eYFP-(control) expressing HEK293 cells was studied in the presence of seven different dicloxacillin or flucloxacillin concentrations and in the absence of inhibitors (vehicle) for 2 min. Rosuvastatin uptake into control cells was subtracted from the uptake into OATP expressing cells for each inhibitor concentration, and the results are presented relative to the uptake in the vehicle groups that were set to 100% transport. Solid lines show the sigmoidal fittings that were used to calculate the half-maximal inhibitory concentrations (IC_50_ values are presented in Table 1). Data points represent mean values of triplicate samples in two independent experiments.

**Table 1.** Half-maximal inhibitory concentrations (IC_50_) for the in vitro transporter inhibition by dicloxacillin and flucloxacillin, and evaluation of potential clinical DDIs by static models. The values in bold are above the threshold values for the risk of in vivo inhibition. A DDI risk evaluation for OATP2B1 and OAT4 is not currently included in the FDA in vitro DDI guidance.^30^

| Transporter | Dicloxacillin |  | Flucloxacillin |  |
| --- | --- | --- | --- | --- |
| | IC<sub>50</sub> (95% CI), μM | Calculated R value | IC<sub>50</sub> (95% CI), μM | Calculated R value |
| OATP1B1 | 3.86<br>(2.61-5.64) | **2.4<sup>a</sup>** | 30.7<br>(14.4-69.8) | **1.5<sup>a</sup>** |
| OATP1B3 | 6.68<br>(3.44-13.34) | **1.8<sup>a</sup>** | 20.7<br>(12.5-35.0) | **1.7<sup>a</sup>** |
| OATP2B1 | 35.5<br>(24.4-51.7) | **1.2<sup>a</sup>**<br>**240<sup>b</sup>** | 64.2<br>(48.6-84.5) | **1.2<sup>a</sup>**<br>**137<sup>b</sup>** |
| OAT3 | 19.5<br>(17.3-21.8) | 0.07<sup>c</sup> | 26.6<br>(23.1-30.6) | **0.1<sup>c</sup>** |
| OAT4 | 7.23<br>(6.18-8.45) | **0.19<sup>d</sup>** | 32.7<br>(28.1-38.0) | **0.08<sup>d</sup>** |
| BCRP | 166<br>(97.6-274) | **51<sup>b</sup>** | 379<br>(227-618) | **23<sup>b</sup>** |
| P-gp | 258<br>(173-374) | **33<sup>b</sup>** | $\geq 833$ | **11<sup>b</sup>** |

<sup>a</sup>R= $1 + (I_{in,max,u}/IC_{50})$ and the FDA cut-off for potential in vivo inhibition is $\geq 1.1$
<sup>b</sup>R= $I_{gut}/IC_{50}$ and the FDA cut-off for potential in vivo inhibition is $\geq 10$
<sup>c</sup>R= $I_{max,u}/IC_{50}$ and the FDA cut-off for potential in vivo inhibition is $\geq 0.1$
<sup>d</sup>R= $I_{max,u}/IC_{50}$ and the FDA cut-off for potential in vivo inhibition is $\geq 0.02$ (the threshold value is defined for Multidrug and toxin extrusion (MATE) proteins that are expressed at the apical side of kidney similar to OAT4)
$I_{in,max}$ is the estimated maximum plasma inhibitor concentration at the inlet to the liver, $I_{max,u}$ is the maximal unbound plasma concentration of the interacting drug at steady state (kidney) and $I_{gut}$ is the intestinal luminal concentration of the interacting drug calculated as the dose/250mL

Dicloxacillin and flucloxacillin inhibited <50% of fluorescein transport into OAT1-expressing HEK293 cells at 200 µM (Figure 2). Similarly, neither compound substantially inhibited OAT2-mediated transport of methotrexate at 200 µM. In contrast, both compounds strongly inhibited fluorescein transport into OAT3-expressing cells and methotrexate transport into OAT4-expressing cells resulting in similar IC_50_-values for OAT3 and OAT4 as for the OATPs (Table 1). A preincubation step was omitted for the OAT inhibition assays since, currently, preincubation has not been reported to potentiate the inhibition of OATs similar to OATPs.^33^ However, we wanted to further investigate whether preincubation potentiates the inhibition of OAT3 by dicloxacillin and flucloxacillin. An assay with 30 min preincubation resulted in the same IC_50_ values (Figure S4) as without the preincubation step (Table 1).

**Figure 2.**
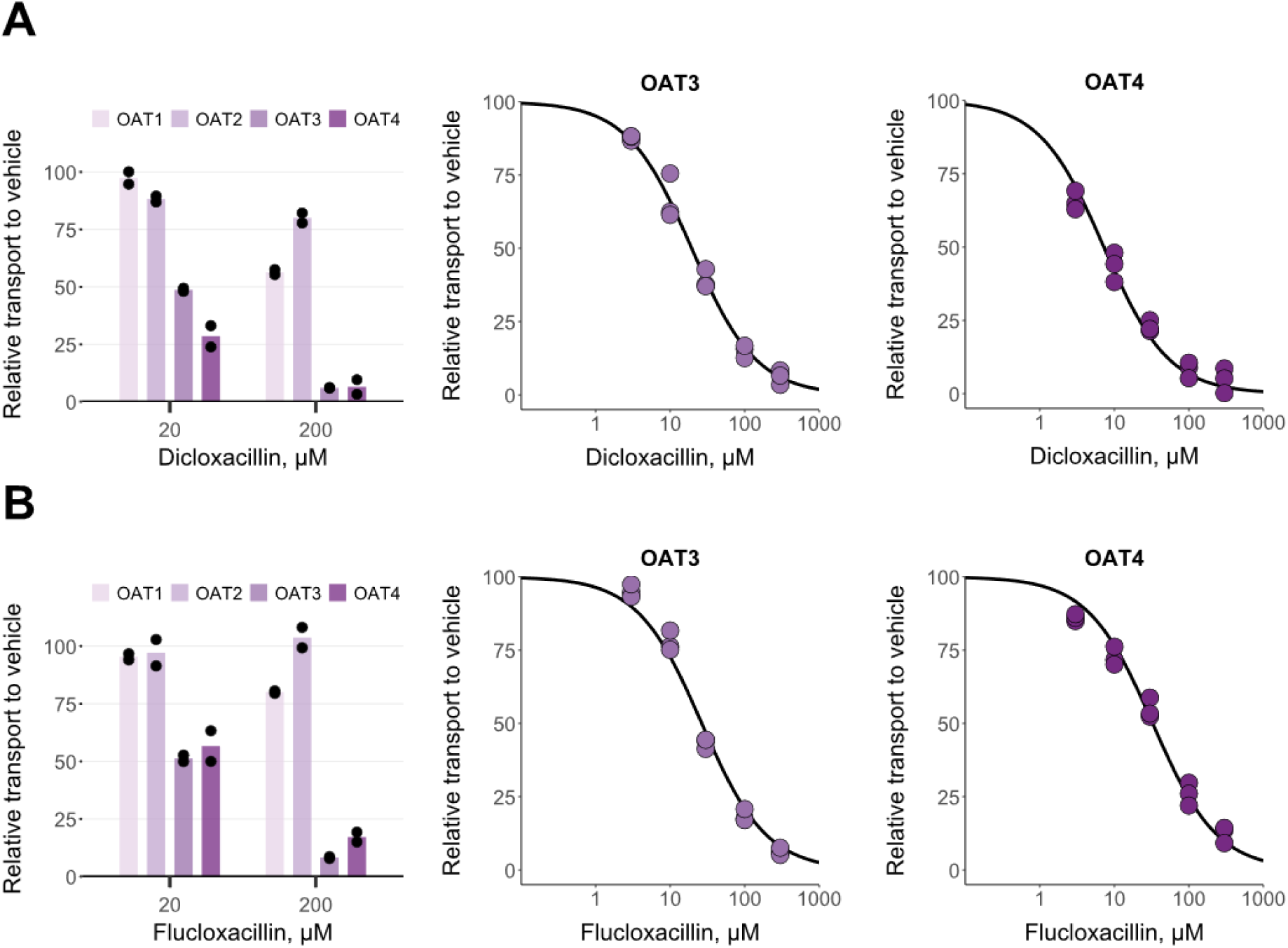
In vitro inhibition of OATs by dicloxacillin (A) and flucloxacillin (B). 5 µM fluorescein transport into OAT1, OAT3 and eYFP (control), and 34 nM methotrexate transport into OAT2-, OAT4- and eYFP-(control) expressing HEK293 cells were studied in the presence of 20 and 200 µM dicloxacillin and flucloxacillin (left side plots) for 5 min (fluorescein) or 10 min (methotrexate). Concentration-dependent inhibition for OAT3 and OAT4 was derived from similar experiments with five different dicloxacillin or flucloxacillin concentrations, including a vehicle sample. For all plots, the uptake into control cells was subtracted from the uptake into OAT-expressing cells for each inhibitor concentration, and the results are presented relative to the uptake in the vehicle groups that were set to 100% transport. Solid lines show the sigmoidal fittings that were used to calculate the half-maximal inhibitory concentrations (IC_50_ values presented in Table 1). Data points represent mean values of duplicate samples in three independent experiments. Bars represent mean values of duplicate samples (presented as points) in an experiment.

Regarding the studied efflux inhibition, dicloxacillin and flucloxacillin inhibited BCRP at high concentrations in a concentration-dependent manner (Figure 3A). For P-gp, the highest concentration of dicloxacillin almost completely inhibited transport, while flucloxacillin reduced P-gp activity by 65% at the highest tested concentration of 2100 µM (Figure 3B). The calculated IC_50_ values were within range of 166 to 379 µM (Table 1).

**Figure 3.**
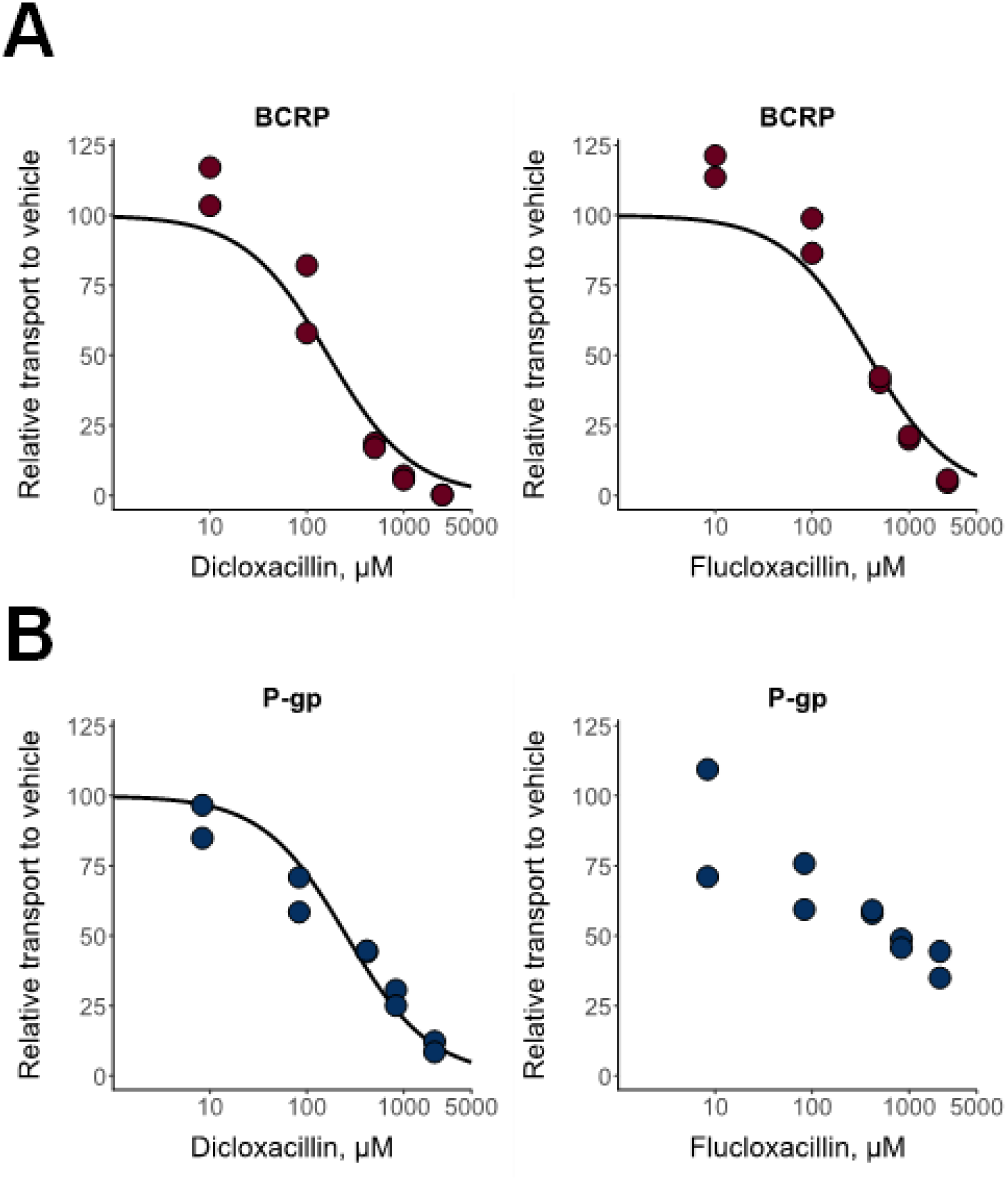
In vitro inhibition of BCRP (A) and P-gp (B) by dicloxacillin and flucloxacillin. BCRP- or P-gp-containing membrane vesicles were incubated with 1 µM rosuvastatin (BCRP) or 1 µM N-methyl-quinidine (P-gp) in the presence of five different concentrations of dicloxacillin and flucloxacillin. Rosuvastatin or N-methyl-quinidine transport in the presence of AMP was subtracted from the uptake in the presence of ATP for each inhibitor concentration, and the results are presented relative to the uptake in the vehicle groups that were set to 100% transport. Solid lines show the sigmoidal fittings that were used to calculate the half-maximal inhibitory concentrations (IC_50_ values presented in Table 1). Data points represent mean values of triplicate samples in two independent experiments.

### Transporter uptake studies

All OATPs and OATs transported dicloxacillin and flucloxacillin, when assayed at 2 and 50 µM concentrations and 10 min transport (Figures 4A-4D). Efflux transporters MRP2, MRP3, MRP4, BCRP and P-gp did not transport dicloxacillin or flucloxacillin in the membrane vesicle assay (Figure S5). We further investigated the time-dependent transport by OATPs and OATs and found that the transport was linear up to 2 min (Figure S6). Because of the low transport activity of OAT1 and OAT2, we did not further characterize these transporters in the transport kinetic assays.

**Figure 4.**
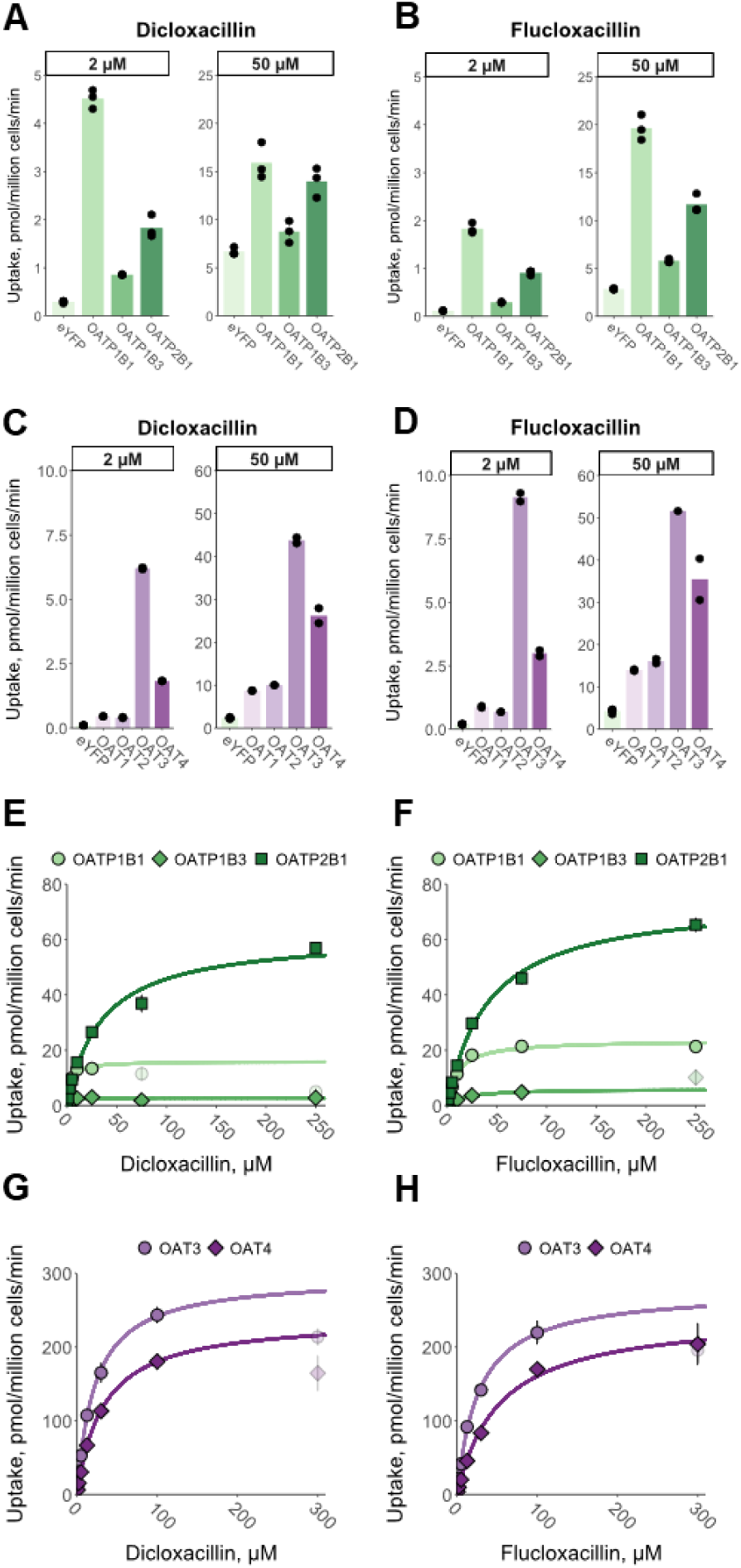
In vitro uptake of dicloxacillin and flucloxacillin into hepatic OATP and renal OAT expressing cells. Transport of 2 µM and 50 µM dicloxacillin and flucloxacillin into OATP1B1-, OATP1B3-, OATP2B1-, OAT1-, OAT2-, OAT3-, OAT4- and eYFP-(control) expressing HEK293 cells were investigated within 10 min incubation (panels A-D). The data points are from one experiment performed with triplicate samples. To study the kinetics for the transport (panels E-H), uptake of seven (OATs) or eight (OATPs) different dicloxacillin and flucloxacillin concentrations into OATP1B1, OATP1B3, OATP2B1, OAT3, OAT4 and eYFP (control) expressing HEK293 cells were studied within 2 min incubation. The uptake into control cells was subtracted from the uptake into transporter expressing cells for each concentration. Because of a substantial deviation from the Michaelis-Menten fittings (solid lines) at the higher concentrations, 75 µM and 250 µM (dicloxacillin, OATP1B1), 250 µM (flucloxacillin, OATP1B3) and 300 µM (dicloxacillin and OATs, flucloxacillin and OAT3) concentrations were excluded from the fittings and are shown with pale colors. Data are presented as mean values of triplicate samples in an experiment (OATPs) or as mean values of duplicate samples in two independent experiments (OATs) with error bars representing standard deviation.

Transport kinetics of dicloxacillin and flucloxacillin by OATP1B1, OATP1B3 and OATP2B1 followed Michaelis-Menten kinetics (Figures 4E and 4F, Table S7). Dicloxacillin and flucloxacillin had the highest affinity for OATP1Bs with K_m_-values of 3 µM (dicloxacillin) and 10-18 µM (flucloxacillin). Similarly, the transport by OAT3 and OAT4 followed Michaelis-Menten kinetics (Figures 4G and 4H, Table S7) but with lower affinities in comparison to OATP1Bs with K_m_-values between 22 µM and 53 µM (Table S7).

### PBPK modelling of transporter inhibition by dicloxacillin and flucloxacillin

To predict the clinical relevance of the observed in vitro inhibition, we calculated the risk for clinical DDIs based on the R-values (Table 1) defined in the FDA in vitro guidance.^30^ Both dicloxacillin and flucloxacillin showed a potential risk for clinical DDIs by inhibiting hepatic OATPs and intestinal BCRP, P-gp and OATP2B1 (Table 1). In addition, the FDA cut-off value was exceeded for the inhibition of OAT4 by dicloxacillin and for the inhibition of OAT3 by flucloxacillin (Table 1). The clinical relevance of OATP, BCRP and P-gp inhibition was further evaluated with PBPK modelling.

The developed PBPK models predicted the clinical pharmacokinetic profiles well for both compounds, with an absolute average fold error of 0.99 and 0.92 for dicloxacillin peak concentration (C_max_) and area under the curve (AUC) and 0.86 and 0.96 for flucloxacillin C_max_ and AUC (Table S8 and S9, Figures S7-S9). In addition, all simulated C_max_ and AUC values were within the calculated drug and study-specific acceptance ranges (Figure S10). Inhibition values derived from the in vitro studies (Table 1) were used as the model inputs for DDI simulations.

Simulations of oral dicloxacillin or flucloxacillin (1000 mg, q 8 h) administered simultaneously with 20 mg rosuvastatin (sensitive OATP and BCRP substrate) predicted only minor changes (≤10%) in rosuvastatin AUC or C_max_ (Table 2, Figures 5A and 5B). Hepatic uptake of rosuvastatin was inhibited up to 27% by dicloxacillin, as was the BCRP-mediated apical efflux clearance in the jejunum (Figure S11A and C). OATP2B1-mediated apical influx of rosuvastatin in the jejunum was predicted to be inhibited up to 62% by dicloxacillin (Figure S11C). On the other hand, flucloxacillin was predicted to have the largest effect on the intestinal apical uptake clearance of rosuvastatin with up to 43% inhibition (Figure S11B and D). In DDI simulations using sensitive P-gp substrates, dabigatran etexilate (150 mg) or digoxin (0.5 mg) as object drugs and dicloxacillin as the precipitant, only minor increases in AUC and C_max_ were predicted by P-gp inhibition (Table 2, Figures 5C and 5D).

**Figure 5.**
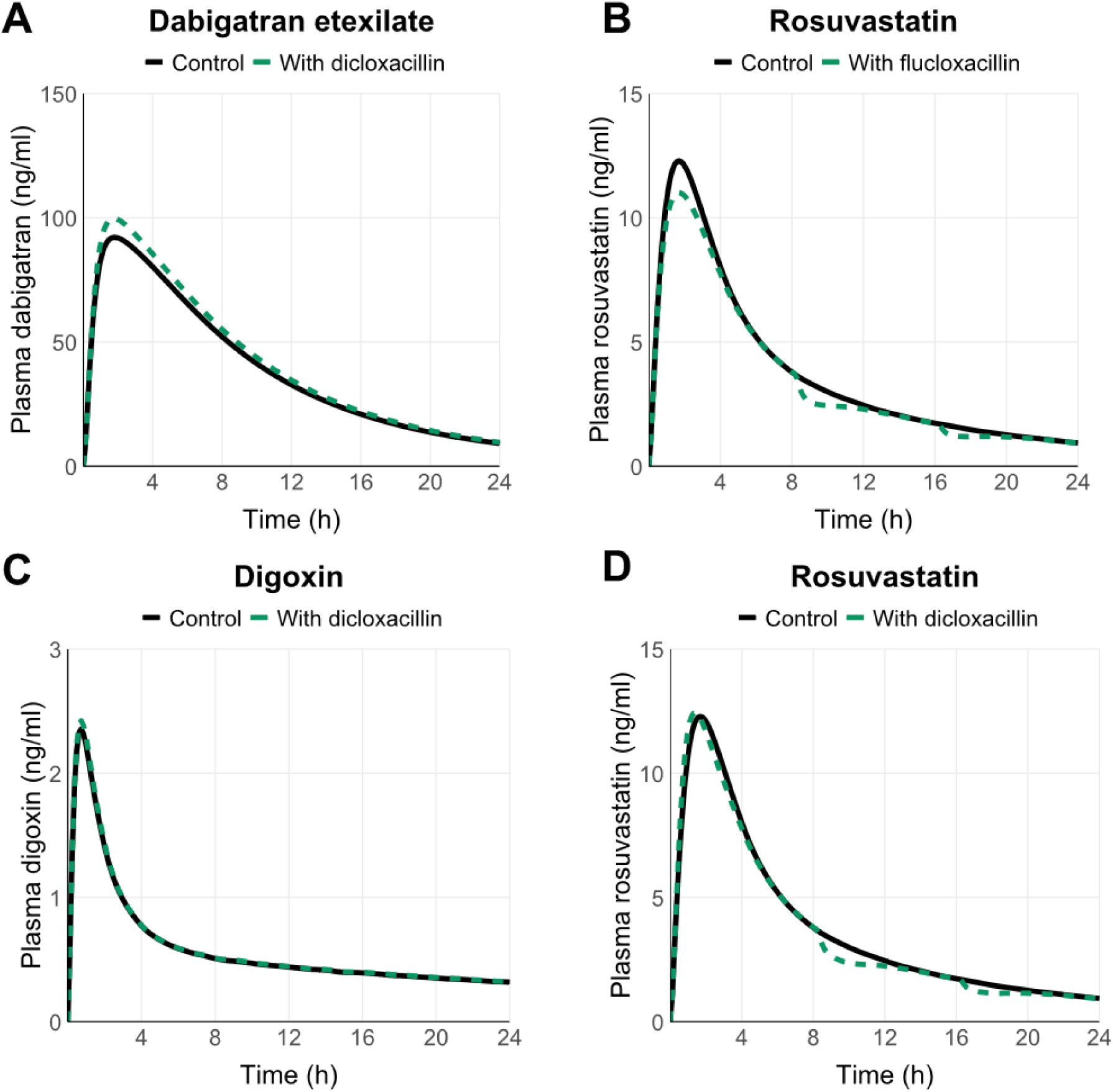
Simulated PK profiles for the object drugs in the DDI simulations in the presence and absence of dicloxacillin (panel A, C and D) or flucloxacillin (panel B). Solid black lines show the predicted mean systemic object drug concentrations in the absence of dicloxacillin or flucloxacillin, while dashed green lines represent the simulations when the object drugs are co-administered together with dicloxacillin or flucloxacillin. The details of DDI simulations and predicted pharmacokinetic parameter changes are reported in Table 2.

**Table 2.** PBPK simulations for DDIs between dicloxacillin and rosuvastatin, dabigatran and digoxin, and between flucloxacillin and rosuvastatin. The PBPK models for the object drugs are built-in library models in Simcyp. DDI simulations were run with 1000 mg per oral dosing of dicloxacillin or flucloxacillin every 8 h. The single dosing of object drugs was either 20 mg rosuvastatin, 150 mg dabigatran etexilate or 0.5 mg digoxin together with the first dose of dicloxacillin or flucloxacillin. The Simcyp Healthy Volunteers population with 10 trials of 10 individuals (20-50 years, 50% female) was employed for the simulations.

|  | **Dicloxacillin** |  |  | **Flucloxacillin** |
| --- | --- | --- | --- | --- |
|  | Rosuvastatin | Dabigatran etexilate<sup>a</sup> | Digoxin | Rosuvastatin |
| AUC ratio | 0.98<br>(0.97 – 1.00) | 1.08<br>(1.07 – 1.08) | 1.02<br>(1.01 – 1.02) | 0.94<br>(0.93 – 0.94) |
| C<sub>max</sub> ratio | 1.10<br>(1.06 – 1.13) | 1.09<br>(1.09 – 1.10) | 1.03<br>(1.02 – 1.03) | 0.94<br>(0.92 – 0.95) |

<sup>a</sup>Pharmacokinetic parameters are for dabigatran, which is the pharmacologically active compound. The AUC and C<sub>max</sub> ratios describe the ratio of the AUC or C<sub>max</sub> values with and without the inhibitor calculated for each virtual study subject. Data are reported as geometric means and 90% confidence intervals.

To examine the effects of dicloxacillin and flucloxacillin on specific OATPs and sinusoidal hepatic uptake clearance in more detail, rosuvastatin DDI simulations were repeated with all the hepatic uptake clearance of rosuvastatin assigned to either OATP1B1, OATP1B3 or OATP2B1. Here, the strongest inhibition (up to 38%) was simulated for dicloxacillin when OATP1B1 accounted for all hepatic rosuvastatin uptake (Figure S11A). This simulation represents a case, where a drug that is taken up into the liver solely by OATP1B1 and has low passive hepatic permeability (1%) is co-administered with dicloxacillin. In this worst-case scenario, the predicted rosuvastatin AUC was almost unchanged, but the C_max_ ratio was 1.24-fold. For flucloxacillin, all models predicted ≤11% changes in the hepatic uptake clearance of rosuvastatin (Figure S11B) and the AUC and C_max_ ratios (0.95 and 0.96-fold) were practically unchanged compared to the simulations with the original model (Table 2).

## DISCUSSION

Dicloxacillin and flucloxacillin have been in clinical use for over 50 years.^5^ Only recently, we reported that both antibiotics cause DDIs by inducing CYP enzymes.^8,10,11,35^ For many older drugs, comprehensive DDI investigations in vitro or in clinical trials were never conducted. Thus, it is unsurprising that we found dicloxacillin and flucloxacillin to inhibit several drug transporters in vitro. We assessed their in vivo DDI potential by employing static model calculations recommended by the FDA (R values)^30^ and further by developing PBPK models to predict hepatic OATP inhibition and intestinal BCRP, P-gp and OATP2B1 inhibition.

The purpose of static models with pre-defined cut-off values for different transporters is to provide a qualitative assessment of potential DDI risk with minimal false negatives.^36,37^ This may, however, result in a high number of false positive signals and higher cut-off values than those recommended by the FDA have been suggested for P-gp and BCRP inhibition ^38–40^ Here, the static model predictions clearly indicated a potential DDI for OATP1Bs and BCRP, whereas the PBPK modelling predicted no significant impact on the systemic exposure of OATP1B and BCRP probe rosuvastatin. The static evaluation likely overestimates the hepatic inlet concentration, which was estimated assuming complete absorption as to depict a worst-case scenario. Unlike R values, mechanistic static modelling, and particularly PBPK modelling, can provide quantitative estimates for the magnitude of possible DDIs. Importantly, as the clearance of drugs is rarely dependent on a single pathway, they allow the simultaneous consideration of multiple pathways and the effect of precipitant drug on each of these pathways.^36,37^ For example, our object drug for the PBPK modeling, rosuvastatin, is transported by multiple transporters in the liver and intestine.^38^ Despite the inhibition of several transporters, the dicloxacillin and flucloxacillin simulations predicted only minor changes in drug exposure (Figure 5, Table 2). This may be explained by the short half-lives of dicloxacillin and flucloxacillin resulting from rapid absorption and clearance as well as their limited accumulation even after multiple dosing (Figure S12).

A novel finding here was that dicloxacillin and flucloxacillin are efficiently transported by OATP1B1, OATP1B3 and OATP2B1. This is unsurprising, as both compounds are anionic carboxylic acids with molecular weights falling within the typical range of OATP substrates.^13^ Still, renal excretion is the major clearance pathway for both antibiotics.^3^ Beringer et al. (2008) found that cyclosporine and probenecid increased the C_max_ and AUC of dicloxacillin slightly (<2-fold).^7^ Furthermore, probenecid decreased the renal clearance of dicloxacillin by 4-fold. Cyclosporine is a strong inhibitor of OATP1B1 and OATP1B3 but does not inhibit OAT1 or OAT3, while probenecid is a strong inhibitor of OAT1 and OAT3 and a weak inhibitor of OATP1B1 and OATP1B3.^41–44^ These in vivo DDI findings can be explained by our results that dicloxacillin is transported by OATP1B1, OATP1B3 and OAT3 and that the inhibition of these transporters affects the pharmacokinetics of dicloxacillin, in particular the renal clearance.

Dicloxacillin and flucloxacillin cause rare but severe idiosyncratic liver injury in humans.^45^ Particularly, the risk of liver injury is well-established for flucloxacillin.^46^ Although the reasons behind this are unknown, transporters may affect the risk of liver toxicity by causing hepatic drug accumulation or, in the case of bile acid transporters, by being inhibited. For example, inhibition of human bile salt export pump (BSEP) may correlate with the increased risk of cholestatic liver toxicity caused by the inhibitor drug.^18,47^ Both dicloxacillin and flucloxacillin inhibit human BSEP.^18,47^ Recently, inhibition of OATP1B1 was reported to be a risk factor for drug-induced cholestasis.^48^ Based on our results, dicloxacillin and flucloxacillin inhibit and are transported by OATP1B1. Interestingly, a genotype of OATP1B1 with reduced activity was not found as a risk factor for liver toxicity caused by flucloxacillin.^49^ Our results may provide new information on the factors that affect the risk of hepatoxicity and liver accumulation of dicloxacillin and flucloxacillin in humans.

Our findings may have implications for other structurally related compounds, such as oxacillin and cloxacillin, which share similar chemical properties with dicloxacillin and flucloxacillin, differing only at two positions of their phenyl rings.^5^ Therefore, it is likely that these two antibiotics may also inhibit human OATPs, OATs and BCRP. Indeed, a study found that cloxacillin inhibits human OAT1, OAT3 and OAT4, and it is transported by OAT3 similarly to what we found here for dicloxacillin and flucloxacillin.^50^ Another study reported similar inhibition of OAT3 by dicloxacillin, cloxacillin and oxacillin.^16^ Furthermore, cloxacillin inhibited OATP1B1 and OATP1B3 transport around 50% at 10 µM in vitro.^15^ The potential transporter DDI liability of oxacillin and cloxacillin should be further investigated.

There are some limitations to our study. We were unable to incorporate the transporter-mediated distribution of dicloxacillin and flucloxacillin into our PBPK models due to the lack of in vitro transporter expression values for scaling and sufficient clinical data for verifying transport pathways. Inclusion of transport may help to better predict intracellular drug exposure, which could have implications for predicting efflux inhibition and liver accumulation. Furthermore, we focused here on short-term inhibition by dicloxacillin and flucloxacillin and, therefore, did not include the previously observed weak P-gp induction^12^ in our dicloxacillin model. However, based on our simulations with dicloxacillin and dabigatran etexilate, induction is expected to have the dominant effect in a clinical scenario, as inhibition without induction was predicted to cause <10% change in dabigatran exposure (Table 2).

In conclusion, dicloxacillin and flucloxacillin inhibit human OATP1B1, OATP1B3, OATP2B1, OAT3, OAT4, BCRP and P-gp in vitro. Static modelling indicated a possible risk for DDIs caused by the inhibition of these transporters, but PBPK simulations predicted no DDI between dicloxacillin or flucloxacillin and the OATP and BCRP substrate rosuvastatin. These findings may be further validated in a clinical study in humans using an appropriate probe substrate or endogenous transporter biomarkers.

## ACKNOWLEDGEMENTS

Certara UK Limited (Simcyp Division) granted access to the Simcyp Simulators through a sponsored academic licence (subject to conditions). The facilities and expertise of the DDCB Unit at the Faculty of Pharmacy, University of Helsinki, supported by HiLIFE and Biocenter Finland, is gratefully acknowledged.

## SUPPLEMENTARY MATERIALS

Supplementary methods and results are reported in Supplementary File

