## SUPPLEMENTARY MATERIALS for "Dicloxacillin and flucloxacillin inhibit hepatic uptake transporters – in vitro investigations and physiologically based pharmacokinetic modelling"

### TABLE OF CONTENTS

| SUPPLEMENTARY MATERIALS AND METHODS | Page |
| --- | --- |
| **Chemicals and reagents** | **3** |
| **In vitro studies** | |
| *Inhibition studies with fluorescent substrates in OATP transduced HEK293 cells* | **3** |
| *Irreversible inhibition assay of OATP1B1* | **3** |
| *Determination of an inhibition constant for the dicloxacillin inhibition of OATP1B1* | **4** |
| *Efflux transport assays in HEK293 membrane vesicles* | **4** |
| **Analytical methods** | **5** |
| **Table S1.** Multiple reaction monitoring parameters for the LC-MS analysis of dicloxacillin, flucloxacillin, rosuvastatin and rosuvastin-$d_6$. | **6** |
| **PBPK modelling** | **6** |
| *Dicloxacillin model* | **7** |
| **Table S2.** Input parameters for the dicloxacillin PBPK model. | **8** |
| *Flucloxacillin model* | **9** |
| **Table S3.** Input parameters for the flucloxacillin PBPK model. | **9** |
| *Dicloxacillin and flucloxacillin simulations* | **11** |
| **Table S4.** Trial parameters used in dicloxacillin simulations. | **11** |
| **Table S5.** Trial parameters | **12** |
| **SUPPLEMENTARY RESULTS** | |
| **In vitro studies** | |
| **Figure S1.** In vitro inhibition of hepatic OATPs by dicloxacillin (a) and flucloxacillin (b) in fluorescent OATP probe substrates assays | **13** |
| **Table S6**. Inhibition constants for the inhibition of transport of OATP1B1, OATP1B3, OATP2B1 by dicloxacillin and flucloxacillin in fluorescent probe assays | **14** |
| **Figure S2**. Reversibility of OATP1B1 inhibition by dicloxacillin and flucloxacillin | **15** |
| **Figure S3.** Determination of the inhibition constant $K_i$ for dicloxacillin and OATP1B1 | **16** |
| **Figure S4.** Effect of a 30 min preincubation step on OAT3 inhibition by dicloxacillin and flucloxacillin | **17** |
| **Figure S5.** Transport of dicloxacillin and flucloxacillin into HEK293 membrane vesicles at 5 and 50 µM | **18** |
| **Figure S6.** Time-dependent transport of dicloxacillin and flucloxacillin into OATP- and OAT-expressing cells | **19** |
| **Table S7.** The transport kinetic parameters for the uptake of dicloxacillin and flucloxacillin into OATP- and OAT-expressing cells | **20** |
| **PBPK modeling** | |
| **Table S8.** $C_{max}$ and AUC values for dicloxacillin PBPK simulations | **21** |
| **Figure S7**. Dicloxacillin simulations for intravenous (IV) and oral (PO) dosing | **22** |
| **Table S9**. $C_{max}$ AUC for flucloxacillin simulations | **23** |
| **Figure S8.** Flucloxacillin simulations for IV dosing | **24** |
| **Figure S9.** Flucloxacillin simulations for PO dosing | **25** |
| **Figure S10.** Comparison of predicted $C_{max}$ and AUC to their acceptance ranges | **26** |
| **Figure S11.** PBPK modelling of sinusoidal hepatic uptake clearance and intestinal apical efflux and uptake clearance | **27** |
| **Figure S12.** Simulated PK profiles for the precipitant drugs in DDI simulations | **28** |
| **SUPPLEMENTARY REFERENCES** | **29** |

#### SUPPLEMENTARY MATERIALS AND METHODS

##### Chemicals and reagents

8- (2- [Fluoresceinyl]aminoethylthio)adenosine- 3', 5'- cyclic monophosphate (8-Fluo-cAMP) was acquired from Biolog Life Science Institute (Bremen, Germany), 4′,5′-dibromofluorescein was acquired from Sigma-Aldrich (St. Louis, MO, USA) and 2′,7′-dichlorofluorescein was acquired from Santa Cruz Biotechnology (Dallas, TX, USA).

##### Inhibition studies with fluorescent substrates in OATP transduced HEK293 cells

The concentrations of fluorescent probe substrates were 1 µM 2′,7′-dichlorofluorescein for OATP1B1, 2.5 µM 8-Fluo-cAMP for OATP1B3 and 1 µM 4′,5′-dibromofluorescein for OATP2B1, and these concentrations were selected to be within or below of previously reported $K_m$-values.[1–3] The transport assays were conducted for 5 min, which is within the previously reported linear ranges of transport for these substrate-transporter combinations.[1,2,4] 8-Fluo-cAMP was dissolved in ultrapure water, while dimethyl sulfoxide (DMSO) was used for all other compounds. The vehicle (DMSO) concentration was 0.3% in all fluorescent substrate assays and the transport assays were conducted as described in the main text and included a 30 min preincubation step. For the quantification of fluorescent substrates, cells were lysed with 120 µl of lysis solution (1% Triton X-100, 5 mM EDTA, 150 mM sodium chloride and 150 mM TRIC-hydrochloride and adjusted to pH 8 with sodium hydroxide) for 30 min at 300 RPM shaking before transferring 80 µl of samples to a transparent 96-well plate containing 20 µl of 0.5 M sodium hydroxide. For each measurement plate, 80 µl of standards, spanning the whole measurement range, were added. The plates were measured with a fluorescent plate reader (Varioskan LUX from ThermoFisher Scientific). The measurement settings were a measurement time of 1000 ms per well, “top”-measurement, excitation bandwidth of 12 nm and with optimized excitation (ex.) and emission (em.) wavelengths for each compound – 495 nm ex. and 525 nm em. for 2′,7′-dichlorofluorescein; 490 nm ex. and 520 nm em. for 8-Fluo-cAMP; 500 nm ex. and 530 nm em. for 4′,5′-dibromofluorescein. Raw values were converted to concentrations, and the mean value of eYFP was subtracted from the mean value of OATP-expressing cells for each sample group. The uptake value into vehicle control sample was set to a value of 100% and all the inhibitor samples are presented relative to the vehicle control sample.

##### Irreversible inhibition assay of OATP1B1

The reversibility of OATP1B1 inhibition by dicloxacillin and flucloxacillin was studied by incubating cells for 15 min in the presence of vehicle or three different inhibitor concentrations. The DMSO (vehicle) concentration was 0.2% in all samples. After the preincubation, solutions were removed and 300 µl of warm TP-buffer was added to each well and incubated for 5 min. The washing was repeated twice, after which the transport assays were conducted in the presence of 1 µM 2′,7′-dichlorofluorescein for 5 min. The wells that were pre-incubated with the inhibitor did not contain any inhibitor during the transport assays, while the wells pre-incubated with the vehicle contained three different inhibitor concentrations. After the transport assays, the samples were treated and analyzed similarly as described in the main text and above. The uptake into vehicle control samples was set to a value of 100% and all the inhibitor samples are presented relative to the vehicle control samples.

**Determination of an inhibition constant for the dicloxacillin inhibition of OATP1B1**

The inhibition constant $K_i$ for a competitive inhibition of OATP1B1 by dicloxacillin was determined in a single experiment in triplicate samples by using 1 µM, 4 µM and 20 µM dicloxacillin and vehicle control (1% DMSO) and varying the concentration of rosuvastatin (1 µM, 2.5 µM, 10 µM, 20 µM, 40 µM and 80 µM). The assays included a 30 min preincubation step with the inhibitor, and the transport assays were conducted for 2 min. Further details are reported in the methods of the main text. The $K_i$ value was calculated by fitting the competitive inhibition model to the data with the nonlinear least squares method (nls) in R programming language environment (R version 4.3.1):

$$Uptake = \frac{V_{max} \times S}{(S + K_m \times (1 + \frac{I}{K_i})}$$

Uptake refers to the rate of uptake in OATP1B1-expressing cells, after subtracting the uptake in control cells, $V_{max}$ is the maximum rate of transport, $K_m$ is the Michaelis constant, S is the concentration of rosuvastatin, I is the concentration of dicloxacillin and $K_i$ is the inhibition constant.

**Efflux transport assays in HEK293 membrane vesicles**

The uptake of dicloxacillin and flucloxacillin into HEK293 membrane vesicles[5] prepared from cells expressing human MRP2, MRP3, MRP4, P-gp, BCRP and eYFP (negative control) was investigated similarly as reported in the main text. Each batch of membrane vesicles was validated with positive controls as reported previously.[5,6] The assays included 5 or 50 µM dicloxacillin and flucloxacillin as the test substrates and were incubated for 10 min. The vehicle (DMSO) concentration was 0.2% in the assays. Each experiment included triplicate samples for both ATP and AMP samples. The elution solvent included dicloxacillin as the internal standard for flucloxacillin samples and flucloxacillin as the internal standard for dicloxacillin samples.

**Analytical methods**

An Acquity UPLC HSS T3 column (1.8 µm particle size, 50x2.1mm from Waters, Milford, MA, USA) equipped with a 0.2 µm in-line filter was employed for the LC-MS analyses and the eluents for chromatographic separation were ultrapure water containing 0.1% formic acid (eluent A) and methanol containing 0.1% formic acid (eluent B). The column and sample chamber temperatures were kept at 40 °C and 15 °C, respectively. The flow rate was 0.4 ml/min, while the sample injection volumes were between 1 and 2 µl and the sample needle was washed with 10% acetonitrile and a mixture of 24.9% ultrapure water, 25% methanol, 25% acetonitrile, 25% isopropanol and 0.1% formic acid after each injection to prevent sample carry-over.

For the chromatographic separation of dicloxacillin and flucloxacillin, the LC gradient was as follows: 55% B (0-0.5 min), 55 to 80% B (0.5-2 min), 95% B (2-2.75 min) and 55% B (2.75-3.25 min). The mass spectrometer was operated in the positive electrospray ionization mode and with a capillary voltage of +2.5kV. Cone and source offset voltages were set to 12 and 60 V, respectively. The source and desolvation temperatures were 150 and 600 °C, respectively. The gas flow settings for cone, desolvation, collision and nebulizer were 150 l/hr, 1000 l/hr, 0.15 ml/min and 7.0 bar, respectively. Dicloxacillin and flucloxacillin were quantified and identified with multiple reaction monitoring (Table S1) and with a dwell time of 50 ms for each transition. The retention times of compounds are presented in Table S1.

For the LC-MS analysis of rosuvastatin and rosuvastatin-$d_6$, the LC gradient was as follows: 55% B (0-0.5 min), 55 to 95% B (0.5-2 min), 95% B (2-3 min) and 55% B (3-3.6 min). The mass spectrometer was operated in the positive electrospray ionization mode and with a capillary voltage of +2.5kV. The mass spectrometer parameters were set to 84 V (cone voltage), 50 V (source offset voltage), 150 °C (source temperature), 600 °C (desolvation temperature), 150 l/hr (cone gas flow), 900 l/hr (desolvation gas flow), 0.15 ml/min (collision gas flow) and 7.0 bar (nebulizer gas flow). Rosuvastatin and rosuvastatin-$d_6$ were quantified and identified with multiple reaction monitoring (Table S1) and with a dwell time of 50 ms for each transition. The retention times are presented in Table S1.

N-methyl-quinidine was analyzed by injecting 5 µl of samples into a Poroshell EC-C18 column (4.6x100mm, 2.7 µm particle size) from Agilent Technologies (Santra Clara, CA, USA) and kept under constant temperature of 40 °C. The sample injection needle was flushed with 10% methanol after each injection to prevent sample carry-over. The chromatographic eluents were ultrapure water with 0.05% phosphoric acid (eluent A) and acetonitrile with 0.05% phosphoric acid (eluent B). The flow rate was 1 ml/min, and the chromatographic method was as follows:

12% B (0-1 min), 12 to 30% B (1-3 min), 90% B (3-4 min) and 12% B (4-6.4 min). N-methyl-quinidine eluted at 2.5 min and was quantified with a fluorescence detector with excitation and emission wavelengths set to 248 nm and 442 nm, respectively.

**Table S1. Multiple reaction monitoring parameters for the LC-MS analysis of dicloxacillin, flucloxacillin, rosuvastatin and rosuvastin-d$_6$.**

| Compound | Retention time, min | Parent, $[H]^+$ m/z | Daughter, m/z | Type of daughter ion | Collision energy, V |
|---|---|---|---|---|---|
| Dicloxacillin | 1.58 | 454.1 | 114.1 | Qualification | 40 |
| Dicloxacillin | | 454.1 | 160.1 | Quantification | 12 |
| Flucloxacillin | 1.38 | 470.0 | 160.1 | Quantification | 12 |
| Flucloxacillin | | 470.0 | 311.1 | Qualification | 16 |
| Rosuvastatin | 1.34 | 482.2 | 258.1 | Quantification | 32 |
| Rosuvastatin | | 482.2 | 300.2 | Qualification | 34 |
| Rosuvastatin-d$_6$ | 1.34 | 488.2 | 264.2 | Quantification | 30 |
| Rosuvastatin-d$_6$ | | 488.2 | 306.2 | Qualification | 38 |

**PBPK modelling**

Full-body PBPK models of dicloxacillin and flucloxacillin were developed in Simcyp version 23 (Certara UK Limited, Sheffield, UK) using published clinical data as described below.

*Clinical data*

WebPlotDigitizer v4 (https://apps.autom eris.io/wpd/) was used to digitize the concentration–time data from the clinical studies. All models were verified using clinical data not used for model development. For dicloxacillin, two studies with intravenous dosing[7,8] and one oral dosing study[9] were used for model development (training set), and five studies[10–14], some of which included several dose levels, were available for model verification (test set). For flucloxacillin, one intravenous study with two dose levels[15] and one study with oral dosing [16] were used for model development (training set), and eight studies[10,14,17–22], some of which included several dose levels or multiple dosing, were available for model verification (test set).

*Model acceptance criteria*

The goodness of fit of simulations was evaluated by comparing predicted and observed concentration-time profiles and calculating peak concentration ($C_{max}$) and area under the concentration-time curve (AUC) ratios ($R_{pred/obs}$) for the drugs using the average absolute fold error (AAFE) calculated as:

$$AAFE = 10^{\frac{1}{n}\Sigma|log(R_{pred/obs})|}$$

where n is the number of studies and $R_{pred/obs}$ refers to predicted versus observed ratios of either $C_{max}$ or AUC. All the datasets used for model verification were included in the AAFE calculation.

In addition, we calculated an acceptance criterion as described by Abduljalil et al. (2014)[23] for PO studies used for verification. The variability of $C_{max}$ and AUC in the clinical studies (σ) was calculated based on the available mean of the coefficient of variation (CV%):

$$\sigma = \sqrt{\ln\left[\left(\frac{CV\%}{100}\right)^2 + 1\right]}$$

Most of the studies did not report a standard deviation (SD) or CV%. For studies reporting interquartile (IQ) range, a quartile coefficient of variation ($CV_Q$%) was calculated using quartiles 1 and 3 (Q1 and Q3) and used for calculating σ:

$$CV_Q = \frac{Q_3 - Q_1}{Q_3 + Q_1}$$

For studies reporting variability as 95% confidence intervals, the SD was back calculated to gain an estimate of CV%.

The σ was then used to calculate upper (A) and lower boundaries (B) for acceptable error for each study (x) using the study mean ($\bar{x}$) and the mean number of participants in the studies (N)[23]:

$$A\bar{x} = exp\left[\ln(\bar{x}) + 4.26\frac{\sigma}{\sqrt{N}}\right]$$

$$B\bar{x} = exp\left[\ln(\bar{x}) - 4.26\frac{\sigma}{\sqrt{N}}\right]$$

Model acceptance was evaluated based on an AAFE between 0.5 and 2 and the containment of the predicted AUC and $C_{max}$ for each study within the calculated acceptance range.

*Dicloxacillin model*

The physicochemical properties of dicloxacillin were sourced from the literature (Table S2). A full-body PBPK model was used to model dicloxacillin distribution. The volume of distribution at steady-state ($V_{ss}$), predicted using the Rodgers and Rowland model (Model 2) within the Simcyp simulator, was close to the previously reported $V_{ss.}$[7,8] The total clearance (CL) of dicloxacillin was calculated as the average from two studies with intravenously (IV) dosed dicloxacillin.[7,8] The renal clearance ($CL_R$) was calculated as the average of available IV and oral (PO) data.[7–9,11,12,14,24] Only studies with 1000 mg dosing or lower were included in the model development and verification due to the possible nonlinear renal clearance of dicloxacillin.[8] The Simcyp retrograde model was used to back calculate the hepatic clearance

($CL_H$) as a human liver microsomal clearance from the in vivo $CL_H$ calculated by subtracting the $CL_R$ from the total CL. The model for oral dosing was optimized using data from the study by Putnam et al. (2005).[9] The absorption rate constant ($k_a$) was calculated from the reported mean absorption time[9] and the fraction absorbed (fa) was optimized manually. The obtained fa of 70% is within the range reported (35 – 76%) for dicloxacillin capsules.[25] Finally, a lag time of 0.3 h was included in the model. The input parameters for the dicloxacillin model are reported in Table S2.

**Table S2. Input parameters for the dicloxacillin PBPK model**.

| Parameter | Value | Reference |
|---|---|---|
| Molecular weight (g/mol) | 470.3 | Pubchem |
| LogPo:w | 2.91 | [26] |
| Compound type | Monoprotic acid | |
| pKa 1 | 2.72 | Mean of values of [26] and [27] |
| PSA (Å$^2$) | 138 | Pubchem |
| HBD | 2 | Pubchem |
| HBA | 7 | Pubchem |
| B:P | 0.65 | Assumed to be the same as for flucloxacillin |
| $f_u$ | 0.032 | Mean of values of [11,28,29] |
| Plasma binding component | HSA | |
| **Absorption model** | | |
| First order | | |
| $f_a$ | 0.7 | Optimized based on [9] |
| $k_a$ ($h^{-1}$) | 1.41 | Calculated from mean absorption time[9] |
| Lag time (h) | 0.3 | Optimized |
| $f_{u,gut}$ | 0.99496 | Predicted |
| $Q_{gut}$ (l/h) | 1.9498 | Predicted |
| $P_{eff,man}$ ($10^{-4}$ cm/s) | 0.23988 | Predicted based on PSA and HBD |
| **Distribution** | | |
| Full PBPK model | | |
| Distribution, $V_{ss}$ (l/kg) | 0.10313 | Predicted, Method 2 (Rodgers and Rowland) |
| $K_p$ values | | |
| Adipose | 0.040866 | |
| Bone | 0.10432 | |
| Brain | 0.058082 | |
| Gut | 0.16726 | |
| Heart | 0.16767 | |
| Kidney | 0.14128 | |
| Liver | 0.096016 | |
| Lung | 0.22225 | |
| Muscle | 0.044361 | |
| Pancreas | 0.070403 | |
| Skin | 0.2893 | |
| Spleen | 0.10794 | |
| **Elimination** | | |
| Total CL (l/h) | 8.885 | Mean value from IV studies[7,8] |
| $CL_H$ (as additional HLM) (µl/min/mg HLM) | 35.426 | Retrograde estimation in Simcyp |
| $CL_R$ (l/h) | 4.71 | Mean value from studies with ≤ 1000 mg dosing[7–9,11,12,14,24] |
| **Interaction** | | |
| *Inhibition* [a] | | |
| OATP1B1 $IC_{50}$ (µM) | 3.9 | This study |
| OATP1B3 $IC_{50}$ (µM) | 6.7 | This study |

| | | |
|---|---|---|
| OATP2B1 $IC_{50}$ (µM) | 35.5 | This study |
| OAT3 $IC_{50}$ (µM) | 19.5 | This study |
| OAT4 $IC_{50}$ (µM) | 7.2 | This study |
| BCRP $IC_{50}$ (µM) | 166 | This study |
| P-gp $IC_{50}$ (µM) | 258 | This study |

[a]Inhibition was included in the intestine, liver and kidney according to the expression of the transporters. BCRP, breast cancer resistance protein; B:P, blood to plasma partition ratio; CL, clearance; $CL_R$, renal clearance; fa, fraction absorbed; fu, fraction unbound; fu,gut, fraction unbound in gut; HBD, number of hydrogen bond donors; HSA, human serum albumin; $CL_{u,int}$, intrinsic unbound clearance; CYP, cytochrome P450; $IC_{50}$, concentration required for 50% of maximal inhibition; $Ind_{50}$, concentration required for 50% of maximal induction; $Ind_{max}$, maximal induction; ka, absorption rate constant; PBPK, physiologically-based pharmacokinetic; OAT; organic anion transporter; OATP, organic anion transporting polypeptide; Peff,man, predicted intestinal permeability in humans; Po:w, oil to water partition coefficient; PSA, polar surface area; Qgut, hybrid parameter describing compound permeability through enterocytes and villous blood flow; $V_{ss}$, volume of distribution at steady-state.

*Flucloxacillin model*

The physicochemical properties of flucloxacillin were sourced from the literature (Table S3). Tissue distribution was described using the full-body PBPK model, with the tissue-to-plasma partition coefficient ($K_p$) predicted based on the Rodgers and Rowland method integrated within the simulator (Method 2). The $K_p$ value for adipose was optimized manually (0.04 to 0.18) so that the predicted $V_{ss}$ matched the mean $V_{ss}$ reported in the literature.[15,17,30] CL values were calculated using a top-down approach from clinical data: The total CL and $CL_R$ were calculated as the mean of values reported in IV studies.[15,17–19,30] $CL_H$ was determined by subtracting $CL_R$ from the total CL. The $CL_H$ was then converted to unbound intrinsic CL ($CL_{u,int,\ H}$) using the well-stirred model, fraction unbound in the plasma and the blood-to-plasma ratio. As flucloxacillin is majorly metabolized by CYP3A4 and CYP2C9, $CL_{u,int}$ for each isoform was calculated from the overall $CL_{u,int,\ H}$. The in vitro intrinsic CL ($CL_{int}$) for each isoform was then calculated using the microsomal protein per gram of liver (32 mg/g) and liver weight (1648 g). This in vitro $CL_{int}$ was further expressed in µl/min/pmol P450 using the enzyme abundance data from the Simcyp Healthy Volunteers population library. The resulting in vitro $CL_{int}$ values were used in the PBPK model. The input parameters for the flucloxacillin model are reported in Table S3.

**Table S3. Input parameters for the flucloxacillin PBPK model.**

| **Parameter** | **Value** | **Reference** |
|---|---|---|
| Molecular weight | 453.9 | Pubchem |
| LogPo:w | 2.6 | [31] |
| Compound type | Monoprotic acid | |
| pKa 1 | 2.76 | [26] |
| PSA | 138 | Pubchem |
| HBD | 2 | Pubchem |
| B:P | 0.65 | [32] |
| $f_u$ | 0.04 | Mean of values in [14,33,34] |
| Plasma binding component | HSA | |

| | | |
|---|---|---|
| **Absorption model** | | |
| First order | | |
| $f_a$ | 0.6 | Predicted |
| $k_a$ ($h^{-1}$) | 1.8 | Optimized (reported range 0.17 -18.2[21]) |
| Lag time (h) | 0.3 | |
| $f_{u,gut}$ | 1 | Predicted |
| $Q_{gut}$ | 1.9498 | Predicted |
| $P_{eff,man}$ ($10^{-4}$ cm/s) | 0.23998 | Predicted based on PSA and HBD |
| **Distribution** | | |
| Full PBPK model | | |
| Distribution, $V_{ss}$ (l/kg) | 0.13006 | Method 2 (Rodgers and Rowland) |
| $K_p$ values | | |
| Adipose | 0.132 | Optimized to recover mean $V_{ss}$ from IV studies[15,17,30] |
| Bone | 0.10538 | |
| Brain | 0.060582 | |
| Gut | 0.16956 | |
| Heart | 0.17033 | |
| Kidney | 0.14409 | |
| Liver | 0.098507 | |
| Lung | 0.22481 | |
| Muscle | 0.046943 | |
| Pancreas | 0.07299 | |
| Skin | 0.29236 | |
| Spleen | 0.11067 | |
| **Elimination** | | |
| Total CL (l/h) | 7.75 | Mean value from IV studies[15,17,18,30,33] |
| $CL_H$ (l/h) | 2.65 | Total CL – $CL_R$ |
| $CL_{u,int,CYP3A4}$ (µl/min/pmol P450) | 0.062 | Retrograde calculation from $CL_H$ (2.65 l/h) |
| $CL_{u,int,CYP2C9}$ (µl/min/pmol P450) | 0.124 | |
| $CL_R$ (l/h) | 5.1 | Mean from reported IV studies[15,17,18,30] |
| **Interaction** | | |
| *Induction* | | |
| CYP3A4 $Ind_{50}$ (µM) | 33.1 | [22] |
| CYP3A4 $Ind_{max}$ | 4.6 | [22] |
| *Inhibition* [a] | | |
| OATP1B1 $IC_{50}$ (µM) | 30.7 | This study |
| OATP1B3 $IC_{50}$ (µM) | 20.7 | This study |
| OATP2B1 $IC_{50}$ (µM) | 64.2 | This study |
| OAT3 $IC_{50}$ (µM) | 27.4 | This study |
| OAT4 $IC_{50}$ (µM) | 32.7 | This study |
| BCRP $IC_{50}$ (µM) | 379 | This study |

[a]Inhibition was included in the intestine, liver and kidney according to the expression of the transporters. BCRP, breast cancer resistance protein; B:P, blood to plasma partition ratio; CL, clearance; $CL_H$, hepatic clearance; $CL_R$, renal clearance; fa, fraction absorbed; fu, fraction unbound; $f_{u,gut}$, fraction unbound in gut; HBD, number of hydrogen bond donors; HSA, human serum albumin; $CL_{u,int}$, intrinsic unbound clearance; CYP, cytochrome P450; $IC_{50}$, concentration required for 50% of maximal inhibition; $Ind_{50}$, concentration required for 50% of maximal induction; $Ind_{max}$, maximal induction; ka, absorption rate constant; PBPK, physiologically-based pharmacokinetic; OAT; organic anion transporter; OATP, organic anion transporting polypeptide; $P_{eff,man}$, predicted intestinal permeability in humans; Po:w, oil to water partition coefficient; PSA, polar surface area; $Q_{gut}$, hybrid parameter describing compound permeability through enterocytes and villous blood flow; $V_{ss}$, volume of distribution at steady-state.

*Dicloxacillin and flucloxacillin simulations*

Trial parameters for the development and verification simulations were chosen to match the clinical study, when available (Table S4 and S5). All simulations were performed using the Simcyp “Healthy Volunteers” population. The verified models were used to simulate interaction studies with BCRP, OATP1B1, OATP1B3 and OATP2B1 substrate rosuvastatin (both dicloxacillin and flucloxacillin) and P-glycoprotein substrates dabigatran etexilate and digoxin (dicloxacillin only). In addition, to further study the inhibitory effect of dicloxacillin and flucloxacillin on specific hepatic OATP transporters, the rosuvastatin model was modified so that the total hepatic uptake clearance was mediated only by one of the OATPs. This was done by summing up the hepatic uptake clearance values of sodium taurocholate co-transporting polypeptide (NTCP), OATP1B1, OATP1B3 and OATP2B1, which mediate rosuvastatin uptake in the model. Further, the total uptake clearance was then assigned to each one of the OATPs in turn and the other hepatic uptake transporters were disabled. DDI simulations were performed with this modified model with the transporter-specific $IC_{50}$ values. The effects of the simulated DDIs were evaluated based on changes in $C_{max}$ and AUC values. In addition, the effects on predicted rosuvastatin apical uptake and apical efflux clearance in the intestine and the apical uptake clearance in the liver were examined by comparing clearance values in the presence and absence of dicloxacillin or flucloxacillin.

**Table S4. Trial parameters used in dicloxacillin simulations.** Parameters were matched as closely as possible to those reported in the original clinical studies.

| Study | Age range (years) | Fraction females | Trials[a] | Dosing |
|---|---|---|---|---|
| **Training** | | | | |
| DeSante et al. 1980 [7] | 21 – 34 | 0 | 20 x 5 | IV bolus 250 mg (2 min) |
| Nauta and Mattie 1976 [8] | 22 – 40 | 0.29 | 20 x 4 | IV bolus 1000 mg (1 min) |
| Putnam et al. 2005 [17] | 21 – 50 | 0.32 | 10 x 17 | PO 1000 mg |
| **Testing** | | | | |
| Sutherland et al. 1970 [10] | 20 – 50[b] | 0.5[b] | 10 x 84 | PO 250 mg |
| Jusko et al. 1975 [11] | 22 – 31 | 0.375 | 20 x 8 | PO 6.25 mg/kg |
| Beringer et al. 2008 [9] | 23 – 28[c] | 0.27 | 10 x 11 | PO 500 mg |
| Gravenkemper et al. 1965 [13] | 20 – 40 | 0.5[b] | 10 x 12 | PO 500 mg |
| Røder et al. 1995 [14] | 32 – 41[c] | 0.5 | 10 x 12 | PO 500 mg |
| Røder et al. 1995 [14] | 32 – 41[c] | 0.5 | 10 x 12 | PO 750 mg |
| Gravenkemper et al. 1965 [13] | 20 – 40 | 0.5[b] | 10 x 12 | PO 1000 mg |
| Røder et al. 1995 [14] | 32 – 41[c] | 0.5 | 10 x 12 | PO 1000 mg |
| **Interaction simulations** | | | | |
| Rosuvastatin | 20 – 50 | 0.5 | 10 x 10 | Rosuvastatin 20 mg (SD, 0 h)<br>Dicloxacillin 1000 mg, q 8 h |
| Dabigatran etexilate | 20 – 50 | 0.5 | 10 x 10 | Dabigatran etexilate 150 mg (SD, 0 h)<br>Dicloxacillin 1000 mg, q 8 h |

| | | | | |
|---|---|---|---|---|
| Digoxin | 20 – 50 | 0.5 | 10 x 10 | Digoxin 0.5 mg (SD, 0 h)<br>Dicloxacillin 1000 mg, q 8 h |

[a]When the number of subjects in the clinical trial was < 10, 20 trials were simulated.
[b]When information was missing, a default range of 20 – 50 years and 50% females was used
[c]Selected age range is based on the reported interquartile range.
IV, intravenous; PO, oral; q 8 h, every 8 h; SD, single dose.

**Table S5. Trial parameters used in flucloxacillin simulations.** Parameters were matched as closely as possible to those reported in the original clinical studies.

| Study | Age range (years) | Fraction females | Trials[a] | Dosing |
|---|---|---|---|---|
| **Training** | | | | |
| Landersdorfer et al. 2007 [15] | 23 – 34 | 0.5 | 10 x 10 | IV infusion (5 min) 500 mg |
| Landersdorfer et al. 2007 [15] | 23 – 34 | 0.5 | 10 x 10 | IV infusion (5 min) 1000 mg |
| Everts et al. 2020 [16] | 18 – 33 | 0.64 | 10 x 11 | PO 1000 mg |
| **Testing** | | | | |
| Landersdorfer et al. 2008 [17] | 18 – 35[b] | 0.5 | 10 x 10 | IV infusion (5 min) 500 mg |
| Adam et al. 1983 [18] | 19 – 34[b] | 0.5 | 10 x 12 | IV infusion (5 min) 1000 mg |
| Landersdorfer et al. 2008 [17] | 18 – 35[b] | 0.5 | 10 x 10 | IV infusion (5 min) 1000 mg |
| Wise et al. 1980 [33] | 20 – 29 | 0 | 20 x 6 | IV bolus (1 min)[c] 1000 mg |
| Sutherland et al. 1970 [10] | 20 – 50 | 0.5[c] | 10 x 11 | PO 125 mg |
| Bodey et al. 1972 [20] | 20 – 53 | 0.5[c] | 20 x 7 | PO 250 mg |
| Sutherland et al. 1970 [10] | 20 – 50[c] | 0.5[c] | 10 x 48 | PO 250 mg |
| Bodey et al. 1972 [20] | 20 – 53 | 0.5[c] | 20 x 7 | PO 500 mg |
| Sutherland et al. 1970 [10] | 20 – 50[c] | 0.5[c] | 10 x 53 | PO 500 mg |
| Røder et al. 1995 [14] | 32 – 41[d] | 0.5 | 10 x 12 | PO 750 mg |
| Gardiner et al. 2018 [21] | 21 – 38 | 0.42 | 10 x 12 | PO 1000 mg |
| Iversen et al. 2023 [22] | 19 – 27 | 0.33 | 10 x 12 | PO 1000 mg |
| Iversen et al. 2023 [22] | 19 – 27 | 0.33 | 10 x 12 | PO 1000 mg q 8 h |
| **Interaction simulations** | | | | |
| Rosuvastatin | 20 - 50 | 0.5 | 10 x 10 | Rosuvastatin 20 mg (SD, 0 h)<br>Flucloxacillin 1000 mg q 8 h |

[a] When the number of subjects in the clinical trial was < 10, 20 trials were simulated.
[b] Calculated based on reported mean +/- 2*SD (for males).
[c] When information was missing, a default range of 20 – 50 years and 50% females was used
[d] Interquartile range
IV, intravenous; PO, oral; q 8 h, every 8 h

#### SUPPLEMENTARY RESULTS

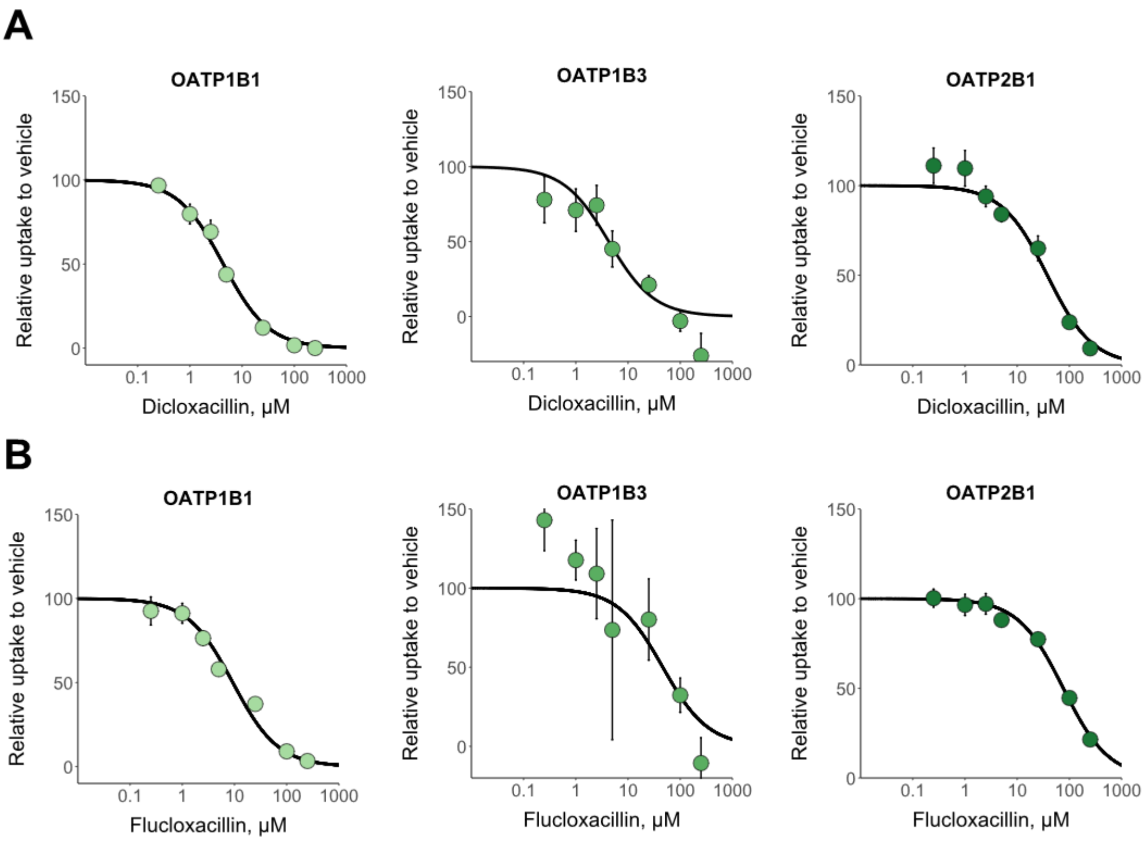

**Figure S1. In vitro inhibition of hepatic OATPs by dicloxacillin (A) and flucloxacillin (B) in fluorescent OATP probe substrates assays.** The fluorescent probe substrates were 1 μM 2′,7′-dichlorofluorescein for OATP1B1, 2.5 μM 8-Fluo-cAMP for OATP1B3 and 1 μM 4′,5′-dibromofluorescein for OATP2B1, and the assays were conducted for 5 min, including a 30 min preincubation step with the inhibitors. The transport into OATP1B1, OATP1B3, OATP2B1- and eYFP (control) -expressing HEK293 cells was studied in the presence of seven different dicloxacillin or flucloxacillin concentrations and in the absence of inhibitors (vehicle). The uptake into control cells was subtracted from the uptake into OATP-expressing cells for each inhibitor concentration, and the results are presented relative to the uptake in the vehicle groups that were set to 100% transport. Solid lines show the sigmoidal fittings that were used to calculate the half-maximal inhibitory concentrations ($IC_{50}$ values are presented in Table S6). Data points represent mean of triplicate samples in an experiment and error bars represent the standard deviation.

**Table S6. In vitro half-maximal inhibitory concentration ($IC_{50}$ value) for the inhibition of transport of OATP1B1, OATP1B3, OATP2B1 by dicloxacillin and flucloxacillin in fluorescent probe assays (Figure S1).**

| | **Dicloxacillin** | **Flucloxacillin** |
|---|---|---|
| **Transporter** | **$IC_{50}$ (95% CI),** μM | **$IC_{50}$ (95% CI),** μM |
| OATP1B1 | 3.59<br>(4.36-5.30) | 9.28<br>(6.31-13.9) |
| OATP1B3 | 4.28<br>(1.66-11.6) | 46.3<br>(12.4-209) |
| OATP2B1 | 37.6<br>(22.4-62.8) | 76.4<br>(63.1-92.4) |

CI, confidence interval

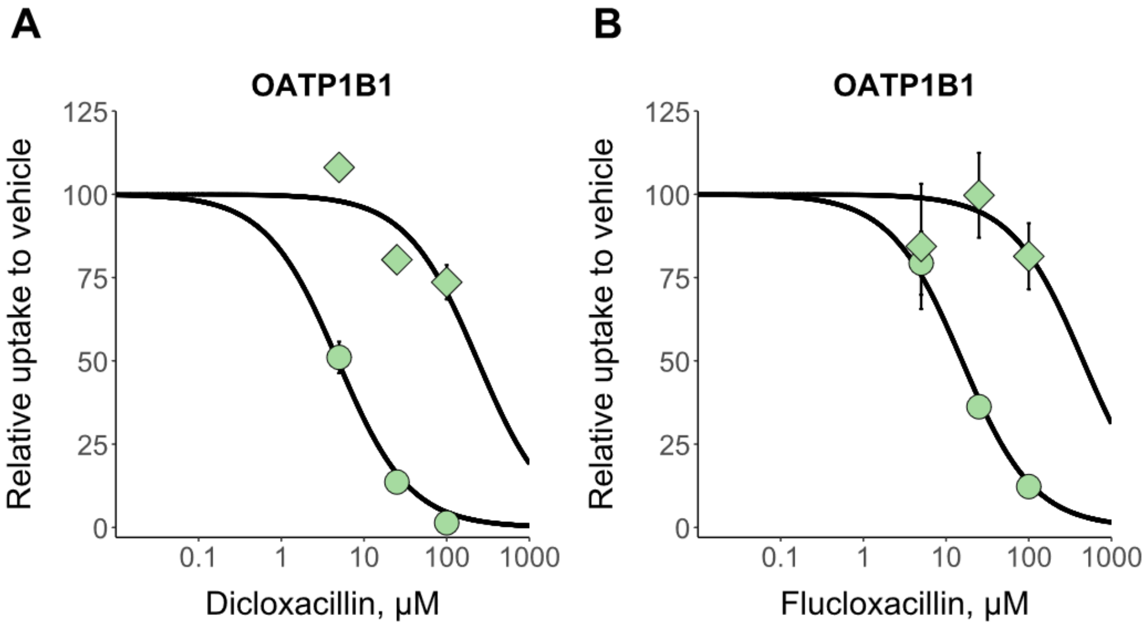

**Figure S2. Reversibility of OATP1B1 inhibition by dicloxacillin (A) and flucloxacillin (B).** The $IC_{50}$-values derived from the fittings for dicloxacillin (A) are 238 µM (95% CIs not available), when the inhibitor was present only during the preincubation after which cells were washed three times before the transport assay, and 4.82 µM (2.93-7.59 µM, 95% CI), when the inhibitor was present only during the transport assay. The $IC_{50}$-values derived from the fittings for flucloxacillin (B) are 455 µM (95% CIs not available), when in the inhibitor was present only during the preincubation after which cells were washed three times before the transport assay, and 15.6 µM (10.4-23.3 µM, 95% CI), when the inhibitor was present only during the transport assay. Each data point represents mean value of triplicate samples in an experiment and error bars represent the standard deviation. Circles represent the samples when inhibitor was present during the transport assays, while squares represent the samples when inhibitor was present only during the preincubation period.

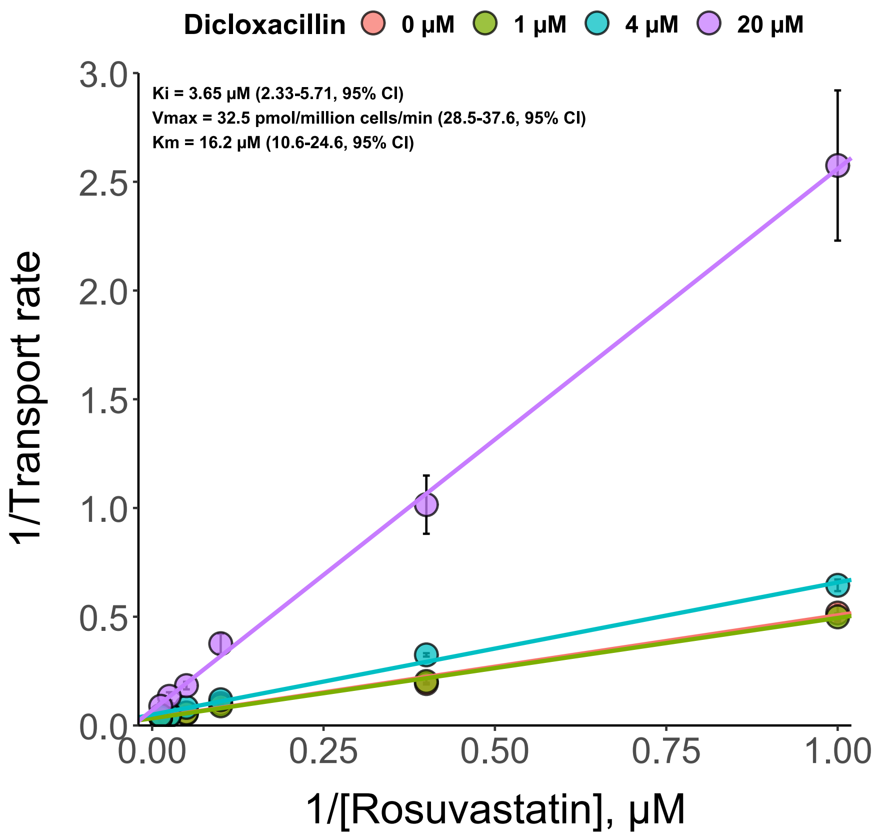

**Figure S3. Determination of the inhibition constant $K_i$ for dicloxacillin and OATP1B1.** Three different dicloxacillin concentrations (1 µM, 4 µM and 20 µM) and a vehicle control (0 µM) were investigated for the inhibition of six different rosuvastatin concentrations (1 µM, 2.5 µM, 10 µM, 20 µM, 40 µM and 80 µM). The Lineweaver-Burk transformation is shown for the data. The uptake rate is derived by subtracting the uptake into control cells from the uptake into OATP1B1-expressing cells. Solid lines show the fittings of the competitive inhibition model to the data. Data is mean of triplicate samples from an experiment and error bars represent standard deviation.

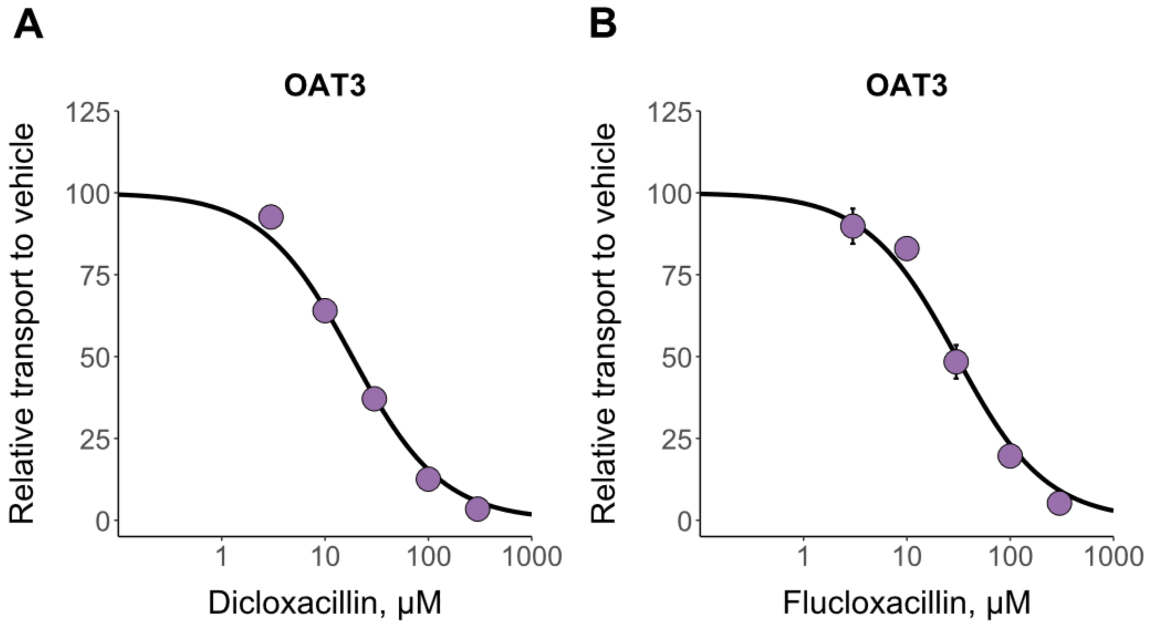

**Figure S4. Effect of a 30 min preincubation step on OAT3 inhibition by dicloxacillin (A) and flucloxacillin (B).** 5 µM fluorescein transport into OAT3- and eYFP- (control) expressing HEK293 cells were studied in the presence five different dicloxacillin or flucloxacillin concentrations, including a vehicle sample. Cells were pre-incubated with the inhibitors and vehicle for 30 min before the assay was initiated with fresh solutions of inhibitors and substrate. The uptake into control cells was subtracted from the uptake into OAT3-expressing cells for each inhibitor concentration, and the results are presented relative to the uptake in the vehicle groups that were set to 100% transport. Solid lines show the sigmoidal fittings that were used to calculate the half-maximal inhibitory concentrations ($IC_{50}$ values). $IC_{50}$-values derived from the fittings are 18.3 µM (13.8-24 µM, 95% CI) and 30.1 µM (21.4-42.5 µM, 95% CI) for dicloxacillin (A) and flucloxacillin (B), respectively. Data points represent mean of duplicate samples from an experiment and error bars represent standard deviation.

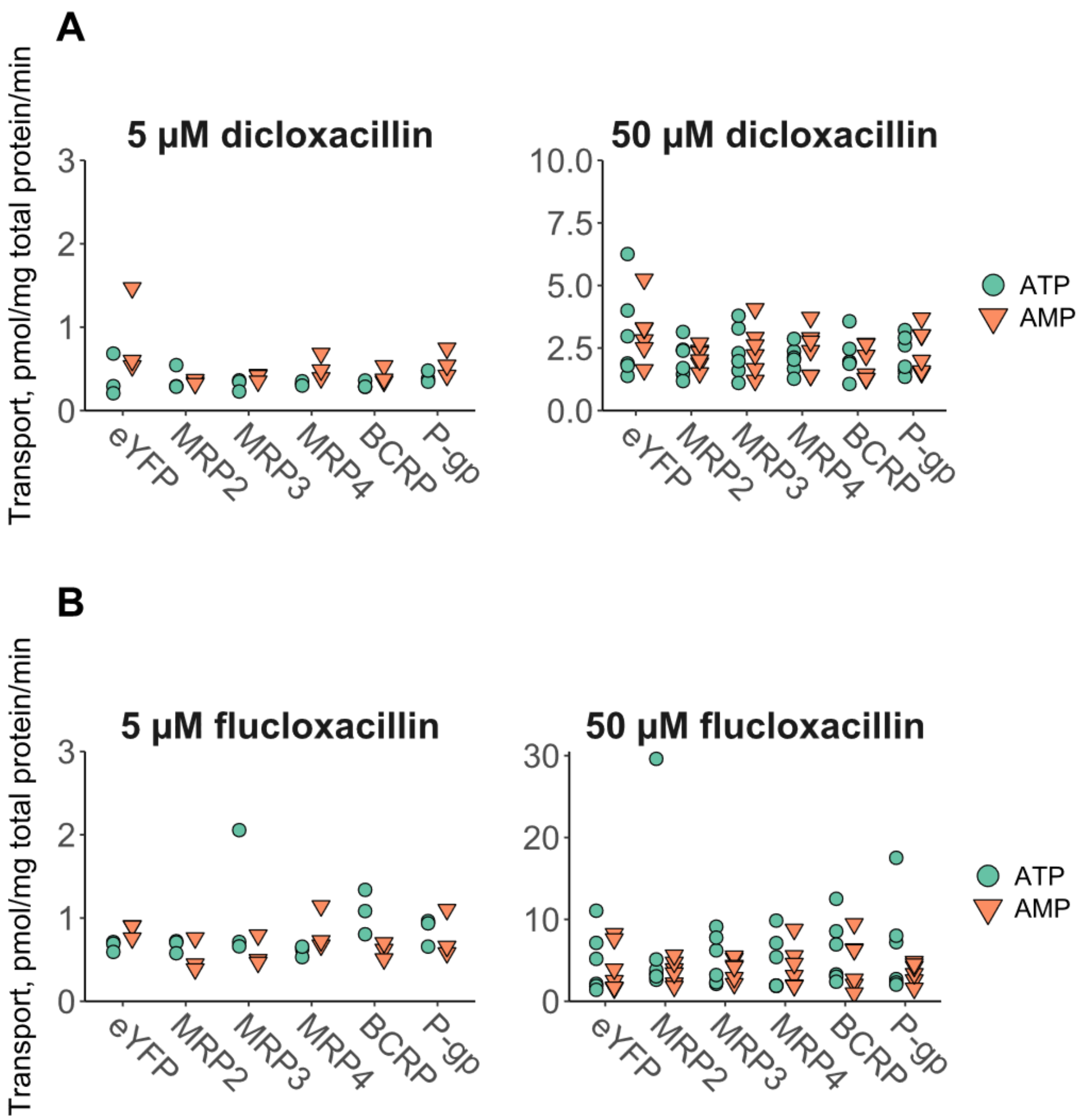

**Figure S5. Transport of dicloxacillin (A) and flucloxacillin (B) into HEK293 membrane vesicles at 5 and 50 µM.** The transport assays were conducted for 10 min. Data points represent individual values from one experiment (5 µM) or two independent experiments (50 µM), each conducted in triplicate samples.

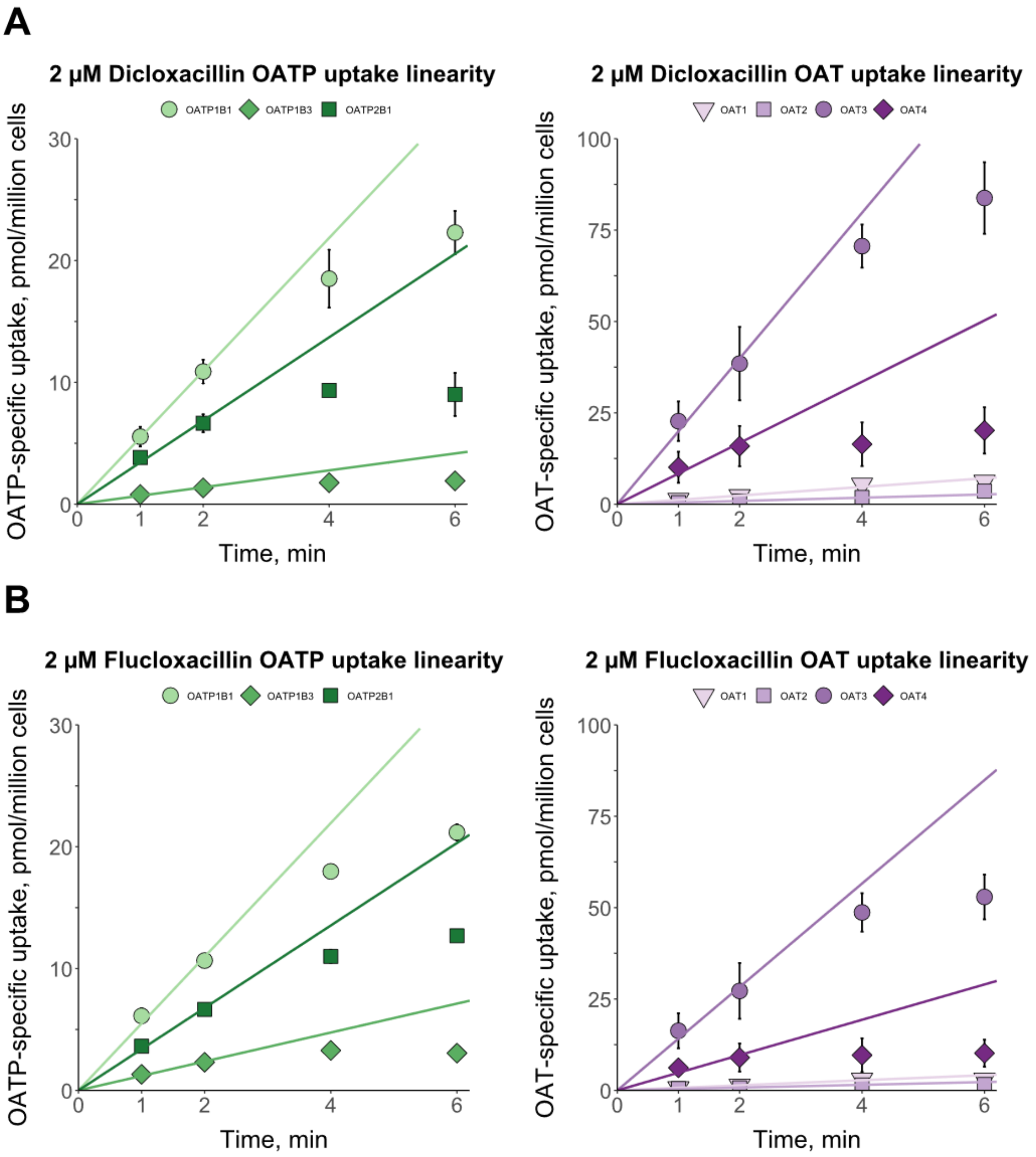

**Figure S6. Time-dependent transport of dicloxacillin (A) and flucloxacillin (B) into OATP- and OAT-expressing cells.** Uptake of 2 µM dicloxacillin and flucloxacillin into OATP1B1-, OATP1B3-, OATP2B1-, OAT1-, OAT2-, OAT3-, OAT4- and eYFP- (control) expressing HEK293 cells were studied within 1, 2, 4 and 6 min. The uptake into control cells was subtracted from the uptake into transporter-expressing cells for each time point. Solid lines show the linear fittings for the transport between 0-, 1- and 2-min. Data are presented as mean values from triplicate samples in an experiment (OATPs) or as mean values from duplicate samples in two experiments (OATs) and error bars represent standard deviation.

**Table S7. The transport kinetic parameters for the uptake of dicloxacillin and flucloxacillin into OATP- and OAT-expressing cells.** The data presented in Figures 4C and 4F were fitted in the Michaelis-Menten equation. 95% confidence intervals of the fitting are presented in parentheses.

| | Dicloxacillin | | Flucloxacillin | |
|---|---|---|---|---|
| **Transporter** | **$K_m$** (µM) | **$V_{max}$** (pmol/$10^6$ cells/min) | **$K_m$** (µM) | **$V_{max}$** (pmol/$10^6$ cells/min) |
| OATP1B1 | 3.22 (1.97-5.13) | 15.9 (13.7-18.6) | 10.4 (7.60-14.2) | 23.5 (21.5-25.7) |
| OATP1B3 | 2.81 (0.718-8.07) | 2.69 (2.02-3.46) | 18.3 (14.5-23.1) | 6.02 (5.50-6.61) |
| OATP2B1 | 34.7 (20.7-59.3) | 61.5 (52.2-73.5) | 41.6 (34.6-50.1) | 74.9 (70.4-79.9) |
| OAT3 | 22.4 (18.3-27.4) | 296 (275-319) | 27.0 (22.5-32.6) | 277 (258-299) |
| OAT4 | 32.9 (30.1-36.1) | 239 (231-249) | 52.5 (40.8-67.9) | 245 (225-267) |

**Table S8. Predicted peak concentrations ($C_{max}$) and area under the concentration-time curve (AUC) values for dicloxacillin PBPK simulations.**

| Study | Dosing | Cmax (mg/l)[a] Pred | Obs | $R_{pred/obs}$ | AUC (mg*h/l)[a] Pred | Obs | $R_{pred/obs}$ |
|---|---|---|---|---|---|---|---|
| **Training** | | | | | | | |
| DeSante et al. 1980 [7] | 250 mg IV | 36.5 | 56.5 | **0.65** | 26.6 | 23.8 | **1.12** |
| Nauta and Mattie 1976 [8] | 1000 mg IV | 234 | NA | **NA** | 113 | 114 | **0.99** |
| Putnam et al. 2005 [9] | 1000 mg PO | 34.0 | 31.4 | **1.08** | 74.5 | 78.0 | **0.96** |
| **Testing** | | | | | | | |
| Sutherland et al. 1970 [10] | 250 mg PO | 8.72 | 11.0 | **0.79** | 16.7 | 19.1 | **0.87** |
| Jusko et al. 1975 [11] | 6.25 mg/kg PO | 15.5 | 18.4 | **0.84** | 33.5 | 33.8 | **0.99** |
| Beringer et al. 2008 [12] | 500 mg PO | 15.8 | 11.1 | **1.42** | 34.1 | 36.6 | **0.93** |
| Gravenkemper et al. 1965 [13] | 500 mg PO | 17.5 | 17.4 | **1.00** | 33.3 | 41.1 | **0.81** |
| Røder et al. 1995 [14] | 500 mg PO | 16.8 | 17.3 | **0.97** | 36.4 | 30.8 | **1.18** |
| Røder et al. 1995 [14] | 750 mg PO | 25.2 | 34.4 | **0.73** | 54.6 | 66.8 | **0.82** |
| Gravenkemper et al. 1965 [13] | 1000 mg PO | 34.9 | 28.9 | **1.21** | 66.5 | 72.4 | **0.92** |
| Røder et al. 1995 [14] | 1000 mg PO | 33.6 | 30.7 | **1.10** | 72.8 | 82.5 | **0.88** |

[a]Output value types (mean, median or geometric mean) of simulations were matched with the type reported in the original clinical study. For AUC, the reported value describes either AUC from the first to the last measurement or to infinity depending on the reported data. If data were not reported in the original studies, it was calculated from the digitized concentration-time curves.
Obs, observed values from clinical studies; Pred, predicted; $R_{pred/obs,}$ ratio of the predicted and observed values.

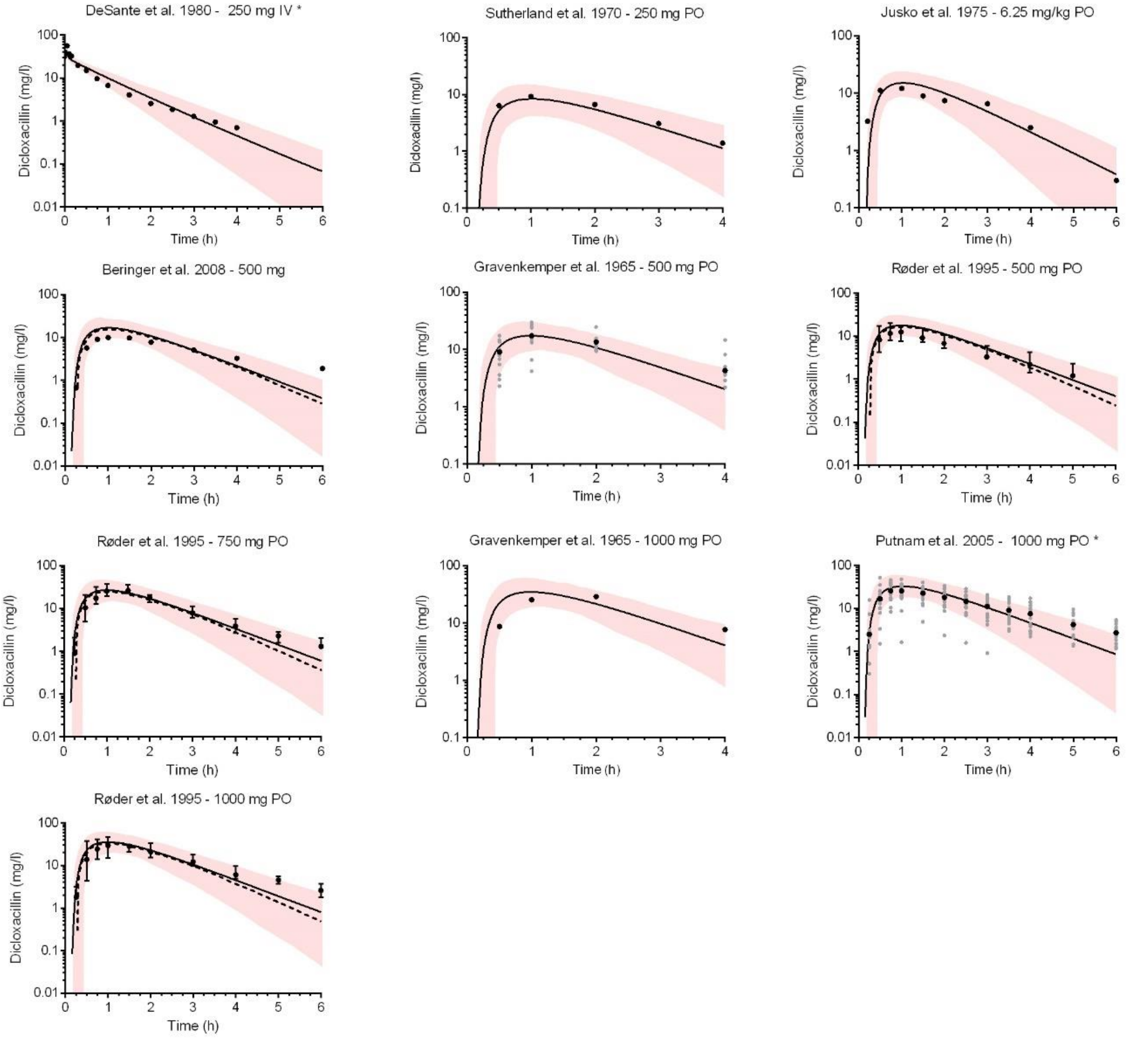

**Figure S7.** Dicloxacillin simulations for intravenous (IV) and oral (PO) dosing. Solid black lines show the predicted mean systemic dicloxacillin concentration and the pink area indicates the area between the 5th and 95th percentiles. Black circles show the mean or median observed data extracted from published clinical studies, whereas grey circles show available individual data. If the observed data was published as median or geometric mean, a dashed line was included in the figure to show the corresponding predicted concentrations. All simulations were performed with trial settings set to match the clinical study in question (Table S4). * Data from DeSante et al.[7] and Putnam et al.[9] were used for the model development. q 6 h, every 6 h.

**Table S9. Predicted peak concentrations ($C_{max}$) and area under the concentration time curve (AUC) values for flucloxacillin simulations.**

| | | Cmax (mg/l)[a] | | | AUC (mg*h/l)[a] | | |
|---|---|---|---|---|---|---|---|
| **Study** | **Dosing** | Pred | Obs | **$R_{pred/obs}$** | Pred | Obs | **$R_{pred/obs}$** |
| **Training** | | | | | | | |
| Landersdorfer et al. 2007 [15] | 500 mg IV | 81.9 | 86.8 | **0.94** | 58.5 | 60.9 | **0.96** |
| Landersdorfer et al. 2007 [15] | 1000 mg IV | 164 | 167 | **0.98** | 117 | 122 | **0.95** |
| Everts et al. 2020 [16] | 1000 mg PO | 25.5 | 31 | **0.82** | 58.1 | 82 | **0.71** |
| **Testing** | | | | | | | |
| Landersdorfer et al. 2008 [17] | 500 mg IV | 82.0 | 87.7 | **0.93** | 54.6 | 54.8 | **1.00** |
| Adam et al. 1983 [18] | 1000 mg IV | 160 | 236 | **0.68** | 125 | 141 | **0.89** |
| Landersdorfer et al. 2008 [17] | 1000 mg IV | 164 | 173 | **0.95** | 109 | 113 | **0.97** |
| Wise et al. 1980 [19] | 1000 mg IV | 94.6 | 140 | **0.68** | 118 | 73.6 | **1.60** |
| Sutherland et al. 1970 [10] | 125 mg PO | 3.46 | 5.8 | **0.60** | 6.09 | 8.28 | **0.74** |
| Bodey et al. 1972 [20] | 250 mg PO | 6.72 | 5.7 | **1.18** | 14.9 | 14.6 | **1.02** |
| Sutherland et al. 1970 [10] | 250 mg PO | 6.55 | 11.0 | **0.60** | 11.6 | 14.4 | **0.81** |
| Bodey et al. 1972 [20] | 500 mg PO | 13.4 | 11.4 | **1.18** | 29.7 | 29.1 | **1.02** |
| Sutherland et al. 1970 [10] | 500 mg PO | 13.0 | 14.7 | **0.89** | 26.6 | 26.6 | **1.00** |
| Røder et al. 1995 [14] | 750 mg PO | 19.4 | 34.5 | **0.56** | 49.2 | 67.8 | **0.73** |
| Gardiner et al. 2018 [21] | 1000 mg PO | 27.3 | 27.4 | **1.00** | 64.9 | 71.6 | **0.91** |
| Iversen et al. 2023 [22] | 1000 mg PO SD | 24.5 | 21 | **1.17** | 60.5 | 51 | **1.19** |
| Iversen et al. 2023 [22] | 1000 mg PO MD (day 9) | 22.7 | 20 | **1.13** | 55.9 | 56 | **1.00** |
| Iversen et al. 2023 [22] | 1000 mg PO MD (day 27) | 22.6 | 24 | **0.94** | 55.7 | 64 | **0.87** |

[a]Output value types (mean, median or geometric mean) of simulations were matched with the type reported in the original clinical study. For AUC, the reported value describes either AUC from the first to the last measurement or to infinity depending on the reported data. If data were not reported in the original studies, it was calculated from the digitized concentration-time curves.
Obs, observed values from clinical studies; Pred, predicted; $R_{pred/obs,}$ ratio of the predicted and observed values.

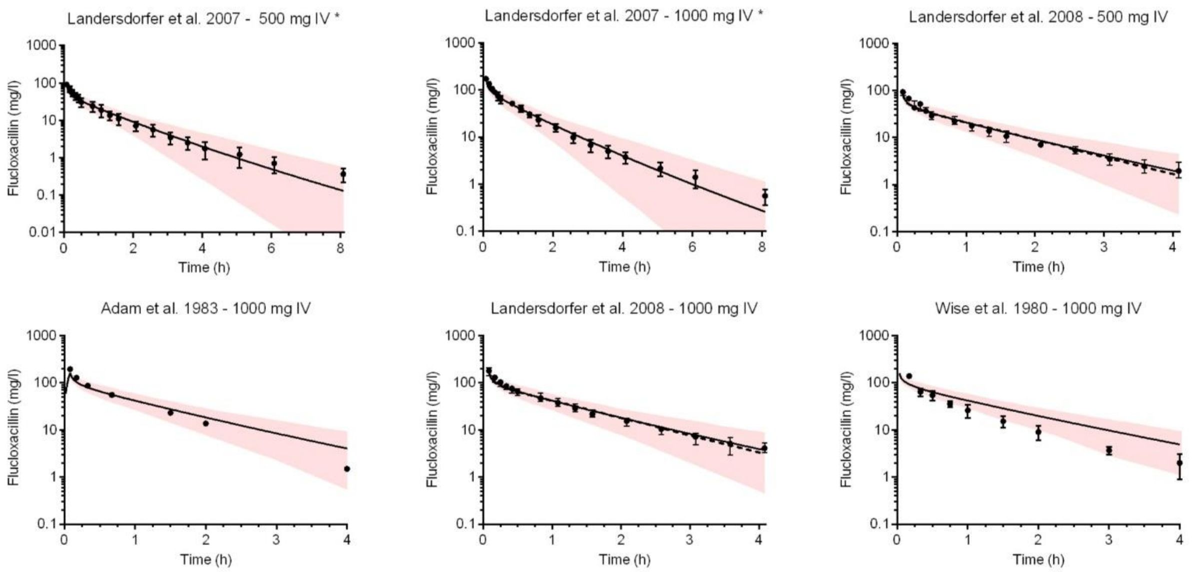

**Figure S8. Flucloxacillin simulations for intravenous (IV) dosing**. Solid black lines show the predicted mean systemic flucloxacillin concentration and the pink area indicates the area between the 5th and 95th percentiles. Black circles show the mean or median observed data extracted from published clinical studies. If the observed data was published as median or geometric mean, a dashed line was included in the figure to show the corresponding simulated concentrations. All simulations were performed with trial settings set to match the clinical study in question (Table S5). *Data from Landersdorfer et al.[15] was used for model development.

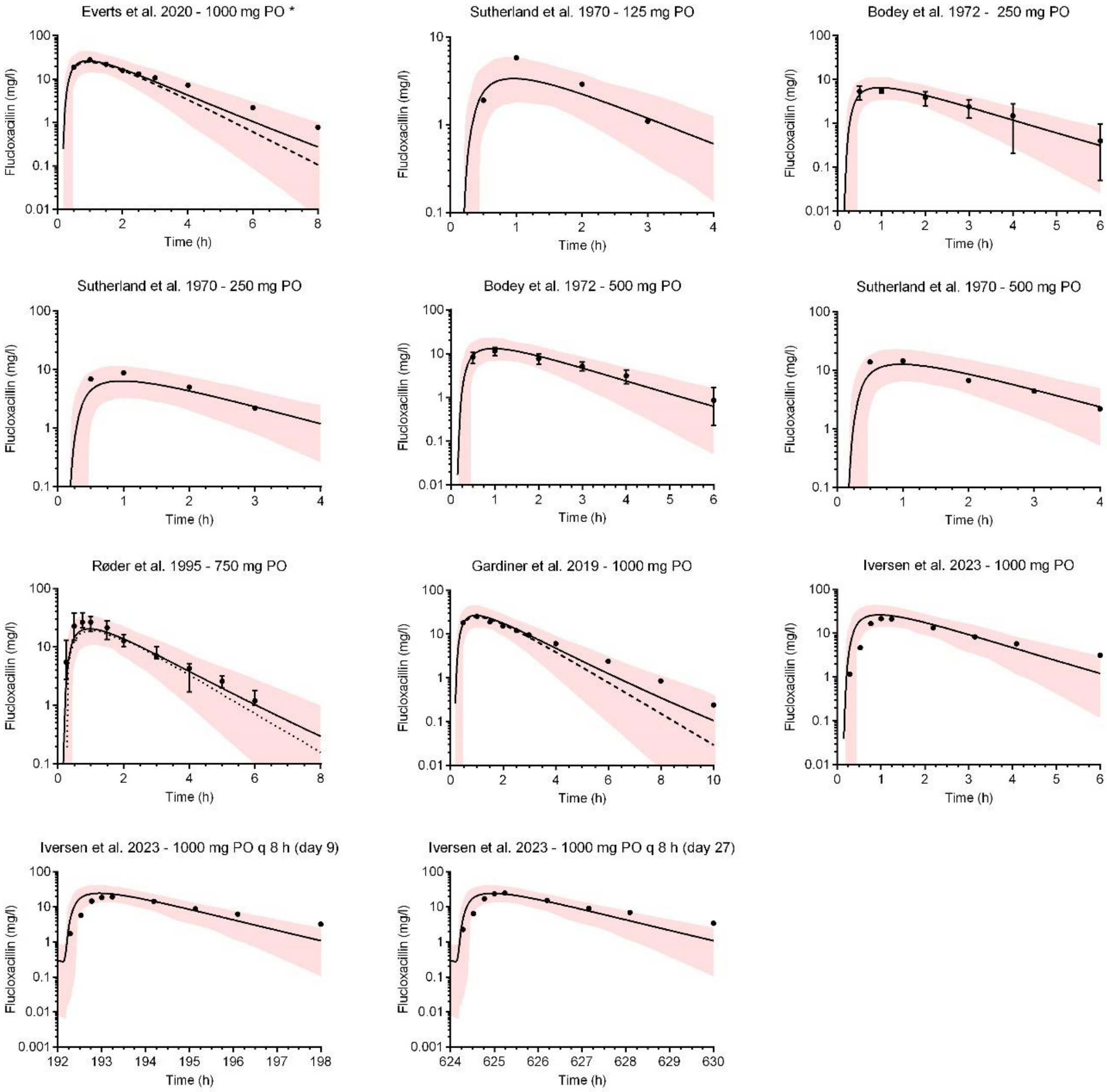

**Figure S9. Flucloxacillin simulations for oral (PO) dosing.** Solid black lines show the predicted mean systemic flucloxacillin concentration and the pink area indicates the area between the 5$^{th}$ and 95$^{th}$ percentiles. Black circles show the mean or median observed data extracted from published clinical studies, whereas red circles show available individual data. If the observed data was published as median or geometric mean, a dashed line was included in the figure to show the corresponding simulated concentrations. All simulations were performed with trial settings set to match the clinical study in question (Table S5). Data from Everts et al.[16] was used for model development. q 8 h, every 8 h.

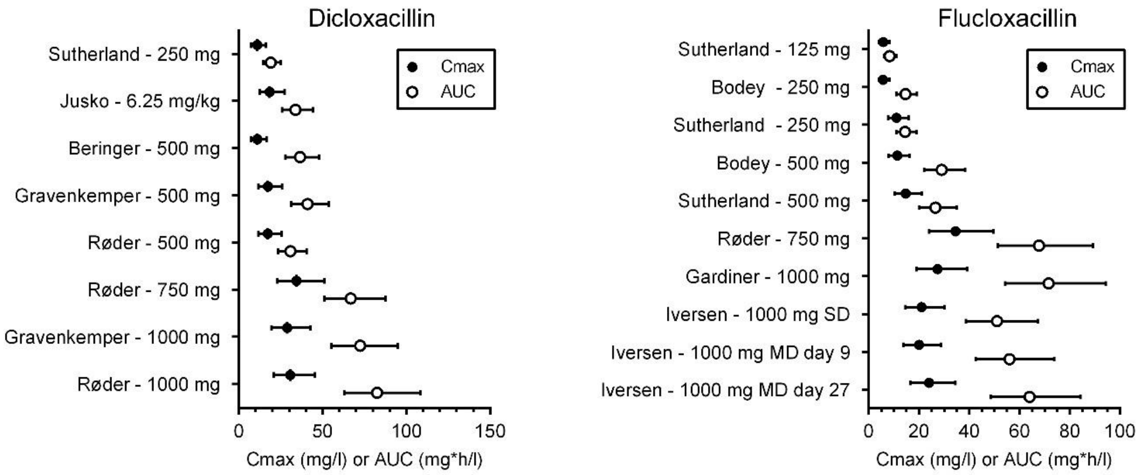

**Figure S10. Comparison of predicted peak concentration ($C_{max}$) and area under the concentration time curve values (AUC) to acceptance ranges of these parameters.** Values for dicloxacillin are presented in the left panel and values for flucloxacillin in the right panel. The circles show the mean predicted $C_{max}$ (closed circles) or AUC (open circles) for each of the simulated clinical studies and the bars represent the span of the acceptance range. The acceptance ranges were calculated according to the method by Abduljalil et al.[23] as described in the *Supplementary materials and methods*. All the predicted mean values are within the estimated acceptance range supporting acceptable model performance. MD, multiple dose; SD, single dose.

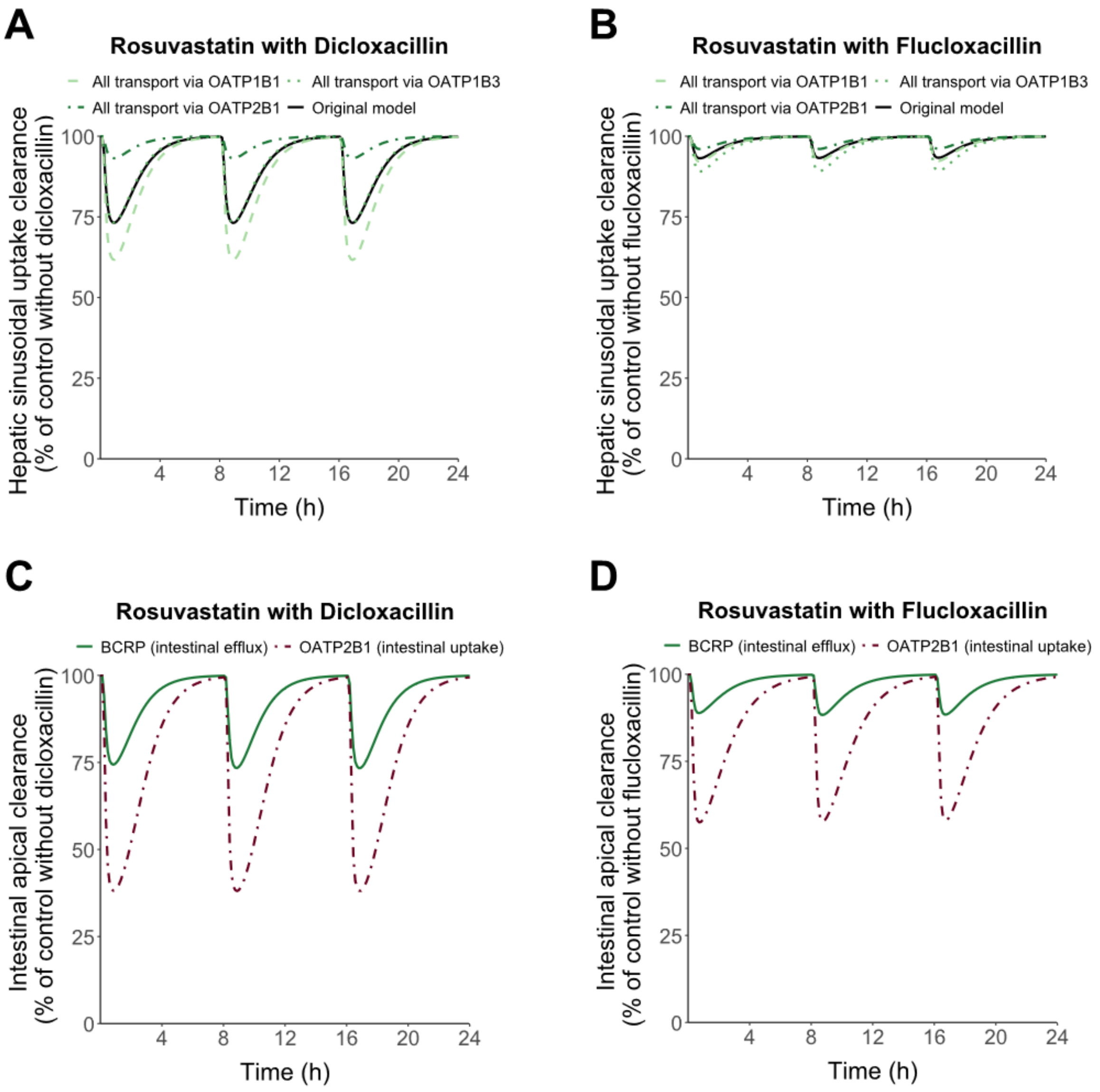

**Figure S11. PBPK modelling of sinusoidal hepatic uptake clearance (A and B) and intestinal apical efflux and uptake clearance (C and D) of rosuvastatin with coadministration of dicloxacillin (A and C) and flucloxacillin (B and D).** The default Simcyp rosuvastatin model was employed with a dose of 20 mg of rosuvastatin. DDI simulations were run with 1000 mg per oral dosing of dicloxacillin or flucloxacillin every 8 h. Hepatic uptake of rosuvastatin is via NTCP, OATP1B1, OATP1B3 and OATP2B1 in the original model and the model was modified by assigning all uptake via either OATP1B1, OATP1B3 or OATP2B1 (panels A and B). The hepatic sinusoidal uptake was compared to a simulation without dicloxacillin or flucloxacillin administration (control), to which data was normalized. The original rosuvastatin model includes intestinal influx by OATP2B1 and intestinal efflux by BCRP at the apical side of jejunum, and the apical clearance values in the presence of inhibitors are presented relative to control without inhibitors.

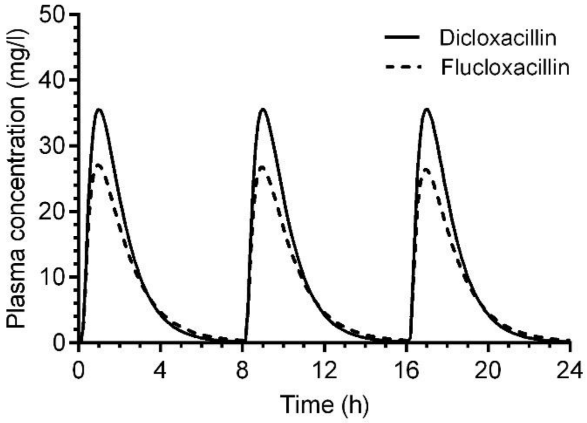

**Figure S12. Simulated mean plasma concentration profiles for the precipitant drugs in the drug-drug interaction (DDI) simulations.** DDI simulations were run with 1000 mg per oral dosing of dicloxacillin (solid line) or flucloxacillin (dashed line) every 8 h. The Simcyp Healthy Volunteers population with 10 trials of 10 individuals (20-50 years, 50% female) was employed for the simulations.
